# Genetic barriers more than ecological adaptations shaped *Serratia marcescens* diversity

**DOI:** 10.1101/2023.07.14.548978

**Authors:** Lodovico Sterzi, Riccardo Nodari, Federico Di Marco, Maria Laura Ferrando, Francesca Saluzzo, Andrea Spitaleri, Hamed Allahverdi, Stella Papaleo, Simona Panelli, Sara Giordana Rimoldi, Gherard Batisti Biffignandi, Marta Corbella, Annalisa Cavallero, Paola Prati, Claudio Farina, Daniela Maria Cirillo, Gianvincenzo Zuccotti, Claudio Bandi, Francesco Comandatore

## Abstract

Bacterial species often comprise well-separated lineages, likely emerged and maintained by genetic isolation and/or ecological divergence. How these two evolutionary actors interact in the shaping of bacterial population structure is currently not fully understood. In this study, we investigated the genetic and ecological drivers underlying the evolution of *Serratia marcescens*, an opportunistic pathogen with high genomic flexibility and able to colonise diverse environments. Comparative genomic analyses revealed a population structure composed of five deeply-demarcated genetic clusters with open pan-genome but limited inter-cluster gene flow, partially explained by Restriction-Modification (R-M) systems incompatibility. Furthermore, a large-scale research on hundred-thousands metagenomic datasets revealed only a partial ecological separation of the clusters. Globally, two clusters only showed a peculiar gene composition and evident ecological adaptations. These results suggest that genetic isolation preceded ecological adaptations in the shaping of the species diversity, suggesting an evolutionary scenario for several bacterial species.

## Introduction

The evolutionary processes shaping the structure of bacterial populations have been deeply investigated and several speciation models have been proposed^1–3^. These models revolve mainly around the two most important mechanisms of genetic variation: mutation and recombination. In 2001, Cohan proposed the “ecotype model” of speciation^4^, which focuses on the role of ecological divergence and selection. In absence of DNA exchange, bacterial lineages periodically accumulate mutations and diverge until one highly adapted lineage emerges and out-competes the other lineages, causing a “clonal sweep” phenomenon. Following this model, stable populations can only exist if they are ecologically diverse enough to avoid competition. A more recent theoretical framework relies on barriers to recombination to explain the origin and maintenance of divergent sequence clusters^5–8^. According to this view, the genetic cohesion is maintained by the persistent genetic exchange between the strains. A speciation event begins when a novel “habitat-specific adaptive allele” spreads within a subpopulation, conferring the ability to occupy a novel ecological niche. The ecological differentiation provides an initial barrier for recombination reducing the genetic exchange among the subpopulations. This process enhances the genetic divergence among these subpopulations, favouring the emergence of genetic barriers and the formation of separate cohesive genotypic clusters. Indeed, recombination rates decrease drastically with sequence divergence^9–11^. This is due to the absence of stretches of identical nucleotides at one or both ends of the recombining DNA sequence, and to the incompatibility between Restriction-Modification (R-M) systems^6,12,13^. The R-M systems are the most widespread bacterial defence systems and rely on a straightforward, efficient mechanism to remove exogenous DNA: a methyltransferase methylates a specific sequence motif on the endogenous DNA and a cognate restriction endonuclease cleaves DNA when the motif is unmethylated. Thus, bacterial populations encoding for noncognate Restriction endonuclease-Methyltransferase (R-M) systems have fewer successful exchanges of genetic material. Ultimately, what emerges is a picture in which bacterial differentiation must be viewed in light of two separate but not exclusive evolutionary drivers: ecology and genetic recombination. More wood has been added to the fire when the concept of “pan-genome” broke into bacterial population genomics. Often, the strains in a bacterial species share only a portion of their gene repertoire, while a consistent part of the genes are owned only by a few strains or lineages (“accessory” genes)^14,15^. Accessory genes could act as a lineage-specific “skill set” with an adaptive impact on the bacterium, involved in the colonisation of a novel ecological niche. Genome-wide speciation models, based on ecological or sexual isolation, are mainly focused on “core” genes^16^ but the analysis of lineage-specific genes can provide pivotal information about the emergence of separated genetic clusters within a species.

*Serratia marcescens* is a Gram-negative opportunistic bacterial pathogen able to cause large outbreaks, in particular in Neonatal Intensive Care Units (NICUs). The bacterium can be also commonly isolated from a multitude of environmental sources, including animal vertebrates^17^, insects^18^, plants^19^, soil^20^ and aquatic environments^21,22^. Several evidence of plant-growth promoting activity^19,23^ further emphasise the versatile lifestyle of this bacterial species. Despite the health concern, only a few large genomic studies about *S. marcescens* are present in literature and the evolution of this species has been poorly investigated so far. During the last years, the first genomic studies about the *S. marcescens* population structure^24–27^ revealed the existence of a certain number of well-defined clades. The most recent and comprehensive studies^26,27^, focused mainly on the distribution of clinically relevant features, proposed the existence of one or more specific hospital-adapted lineages, harbouring antibiotic resistance and/or virulence markers. Moreover, a recent wide genomic study on the whole *Serratia* genus^28^ has highlighted numerous events of niche specialisation associated with specific gene composition, suggesting that the strong ecological plasticity in the genus is fostered by events of gene gain and loss. Although these studies are progressively shedding light on the population structure and main genomic features of *S. marcescens*, many facets about which mechanisms have played a role in the origin and maintenance of this genetic diversity are still unclear.

The aim of this study was to characterise the diversity within *S. marcescens* and to trace signals of how ecology and gene flow affect the population structure of this wide-spread, ubiquitary and versatile bacterial species.

## Results

### Reconstruction of the study Global genomic dataset

The forces shaping the evolution of *Serratia marcescens* were investigated on a large and cured high-quality genomic dataset (labelled “Global dataset”) including 902 genome assemblies. The genomes were selected from a preliminary collection of 1,113 genomes (see Methods, Table S1 for details)). The Global dataset comprises: i) 230 *S. marcescens* genomes sequenced as part of a large study involving six hospitals in Northern Italy (Alvaro et al. 2023, submitted); ii) five additional strains from the same collection of isolates (sequenced ex novo); iii) 667 genomes from public databases. Overall, this is one of the widest genomic datasets analysed in a comparative genomic study on *S. marcescens* so far. Genomes were manually classified into three categories on the basis of the isolation source: 715 clinical, 122 environmental and 29 animal. For 36 strains it was not possible to obtain a reliable classification due to the incompleteness of the related metadata. It must be noted that, as in most studies involving opportunistic pathogens, strains from clinical samples are overrepresented in the dataset.

### The *Serratia marcescens* population structure reveals five phylogenetic clusters

In the first step of the study, the population structure of *S. marcescens* was investigated by combining core Single Nucleotide Polymorphisms (SNP)-based phylogenetic analysis with Principal Coordinates Analysis (PCoA) clustering on coreSNPs and Mash distances. The SNP calling procedure returned a total of 22,290 coreSNPs and the relative rooted Maximum Likelihood (ML) tree is shown in Figure 1a. The unsupervised K-means clustering performed on patristic distances, coreSNPs or Mash distances converged in dividing the *S. marcescens* population in five well-distinguished clusters (Figures S1). The clusters are coherent with the phylogenetic clades (Figure 1a) and demarcated by deep divisions in the tree. Despite Cluster 1 comprises 53% (475/902) of the strains within the Global genomic dataset, the distribution of Average Nucleotide Identity (ANI) between strains of the same cluster^29^, shows that Cluster 4 and Cluster 5 contain clearly more genetic variability than the other clusters (Figure 1b). Interestingly, ANI among strains of different clusters draws near (and in some cases exceeds) the 95% ANI-based species boundary^30^. Indeed, the maximum ANI between the clusters ranges from 96.78% for the Cluster 1 - Cluster 4 pair to the 95.56% for Cluster 3 - Cluster 4 (Figure 1c). Overall, the population structure of *S. marcescens* reveals the existence of well-differentiated genetic clusters with clear genetic boundaries, suggesting a remarkable intraspecies genetic diversity.

**Figure 1.**
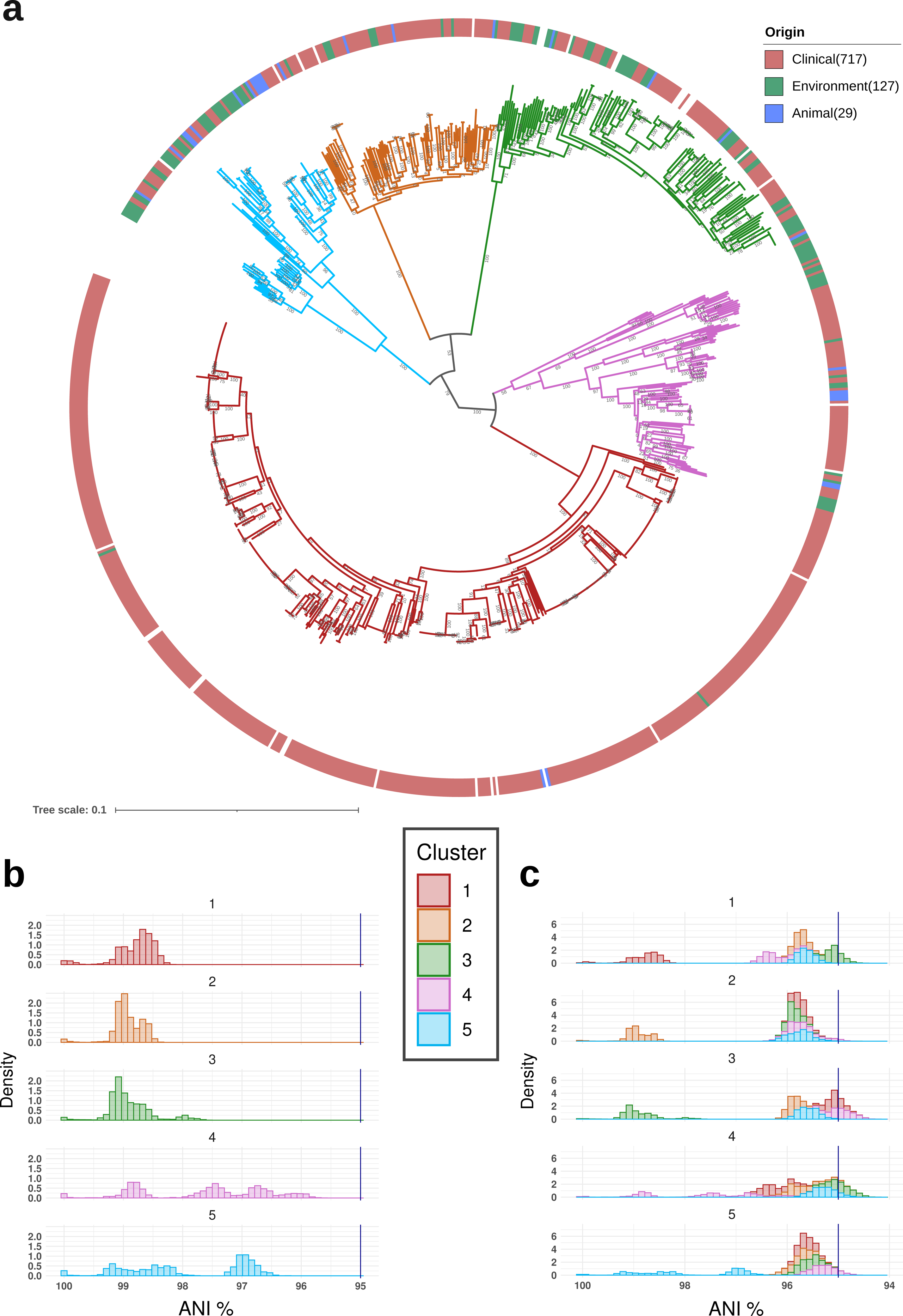
The population structure of *Serratia marcescens*. a) SNP-based Maximum Likelihood (ML) phylogenetic tree of the 902 *Serratia marcescens* strains of the Global genomic dataset. The tree branches’ colours and the inner circle around the tree indicate the five clusters coherently and independently determined applying K-means clustering on patristic distances, coreSNP distances and Mash distances. The circle around the tree indicates the strain isolation source (blank if not traceable from the metadata). Bootstrap values are shown on the tree nodes. b) Distribution of Average Nucleotide Identity (ANI) between *S. marcescens* strains *within* each cluster. The dark blue vertical lines indicate the species identity threshold (95% ANI). c) Distribution of Average Nucleotide Identity (ANI) *within and between* clusters. The dark blue vertical lines indicate the species identity threshold (95% ANI).

### Specific genomic features highlight diversity between clusters

Genomic features, such as genome size and GC content, were compared between the five clusters (Figure S2). Genome size ranges from 4,955,525 bp to 5,896,859 bp and Cluster 4 has a wider genome size in comparison to Cluster 1, Cluster 2, Cluster 3 and Cluster 5. Cluster 1 genomes are also significantly larger than genomes in Cluster 3. Despite all *S. marcescens* strains displaying a percentage of GC content between 58.9% and 60.2%, comparison between clusters showed that Cluster 1 has a markedly higher GC content than Cluster 2, Cluster 3, Cluster 4 and Cluster 5. At the same time, Cluster 2 also has a lower GC content than Cluster 3, Cluster 4 and Cluster 5. These results are coherent with those recently published by Ono and colleagues^26^, in which the authors proposed that the difference in GC content could be a consequence of Horizontal Gene Transfer (HGT) with different donor bacteria. P values of significant combinations are shown in Figure S2 and in Supplementary Material 1.

The synteny analysis performed on 65 complete genomes highlights occasional translocations and inversions occurring among strains of the same Cluster, but synteny is overall preserved in the global population and all clusters share highly syntenic blocks (Figure S3).

Overall, the observed inter-cluster variations in genome size and GC content are coherent with the cluster’s genetic separation described above.

### The *Serratia marcescens* clusters are enriched in specific isolation sources

As expected for a human-associated wide-spread bacterium, all clusters are dispersed in every continent apart from Africa and Oceania, greatly underrepresented in the dataset. However, Chi-squared test has revealed an uneven distribution of the clusters in the global continents (X-squared = 103.11, df = 20, p-value = 3.486e-13) and the analysis of the residuals showed that Cluster 5 is associated with North America and negatively associated with Europe (Figure S4). Moreover, a focus on the spatio-temporal distribution of the 235 strains sampled from six Italian hospitals showed that multiple clusters often coexist within the same hospital in the same time period (Figure S5).

The geographically-balanced analysis of the association between Cluster and isolation source (see Methods) indicates that Cluster 1 is significantly associated with clinical settings and negatively associated with environmental sources (Figure 2). Despite not reaching statistical significance, Cluster 3 and Cluster 5 also display a clear pattern of enrichment in environmental (Cluster 3 and Cluster 5) and animal sources (Cluster 5).

**Figure 2.**
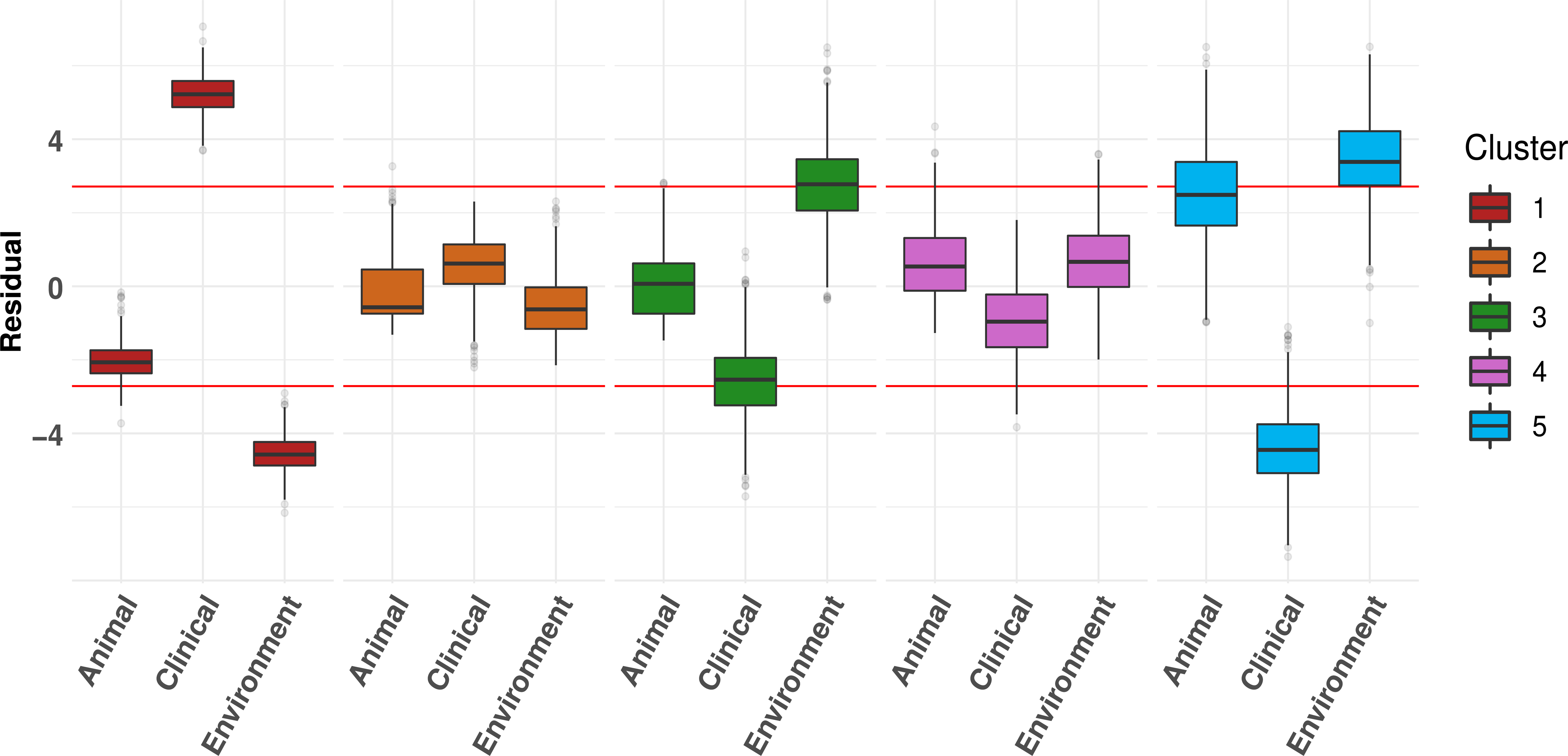
Association between phylogenetic cluster and isolation source. c) Association of *S. marcescens* clusters with animal, clinical or environmental isolation sources. To avoid potential biases due to sampling proximity, Chi-squared test was repeated 1,000 times on geographically-balanced subsets. The box plot illustrates the distribution of the Pearson residuals for each cluster-isolation source combination, and the red horizontal lines demarcate the thresholds for statistical significance of the residual (see Methods): Cluster 1 is associated with clinical samples (>95% of subsets are significant). Cluster 3 is enriched in environmental isolation sources (52% of subsets), and Cluster 5 is enriched in both environmental (76% of subsets) and animal (42% of subsets).

In 2018, Abreo and Altier^24^ proposed that *S. marcescens* could be differentiated in an “environmental” clade and a “clinical” clade. Following studies on larger genomic datasets have refined this idea, suggesting the existence of one or more clinical/hospital-based lineages^26,27^). Our results highlight that certain clusters are enriched in environmental, animal or clinical samples, thus providing a signal of possible ecological specificity of the *S. marcescens* clusters. At the same time, different clusters were frequently isolated from the same hospital in the same period, strongly suggesting that the observed genetic separation cannot be explained only by habitat segregation.

### Two *Serratia marcescens* clusters have highly peculiar gene repertoires

The *S. marcescens* pan--genome comprises a total of 57,700 genes: 2,811 core genes (present in >=95% strains), 3,286 shell genes (>=15% and <95%) and 51,603 cloud genes (<15% of the strains). The pan-genomes of *S. marcescens* and of each single cluster are open (slope of the log-log cumulative curve linear regression < 1, p value < 0.05, Figure S6-7). The five clusters show pan-genomes of different size: Cluster 4 exhibits the largest pan--genome and boosts the species total pan--genome, followed by Cluster 5, Cluster 2, Cluster 3 and lastly Cluster 1. Often, the size of a bacterial pan-genome is considered to be related to the lifestyle of the species, and open pan-genomes are associated with ubiquitary bacterial species with wide ecological niches and high rates of horizontal gene transfer^31^. As shown in Figure 3a, the intensity of gene gain/loss mapped on the phylogenetic tree shows that Cluster 1, Cluster 2 and Cluster 3 exhibit an extensive gene gain/loss on their basal node. Interestingly, major gene gain/loss is also frequent within smaller lineages, reinforcing the assumption that *S. marcescens* undergoes frequent gene turnover.

**Figure 3.**
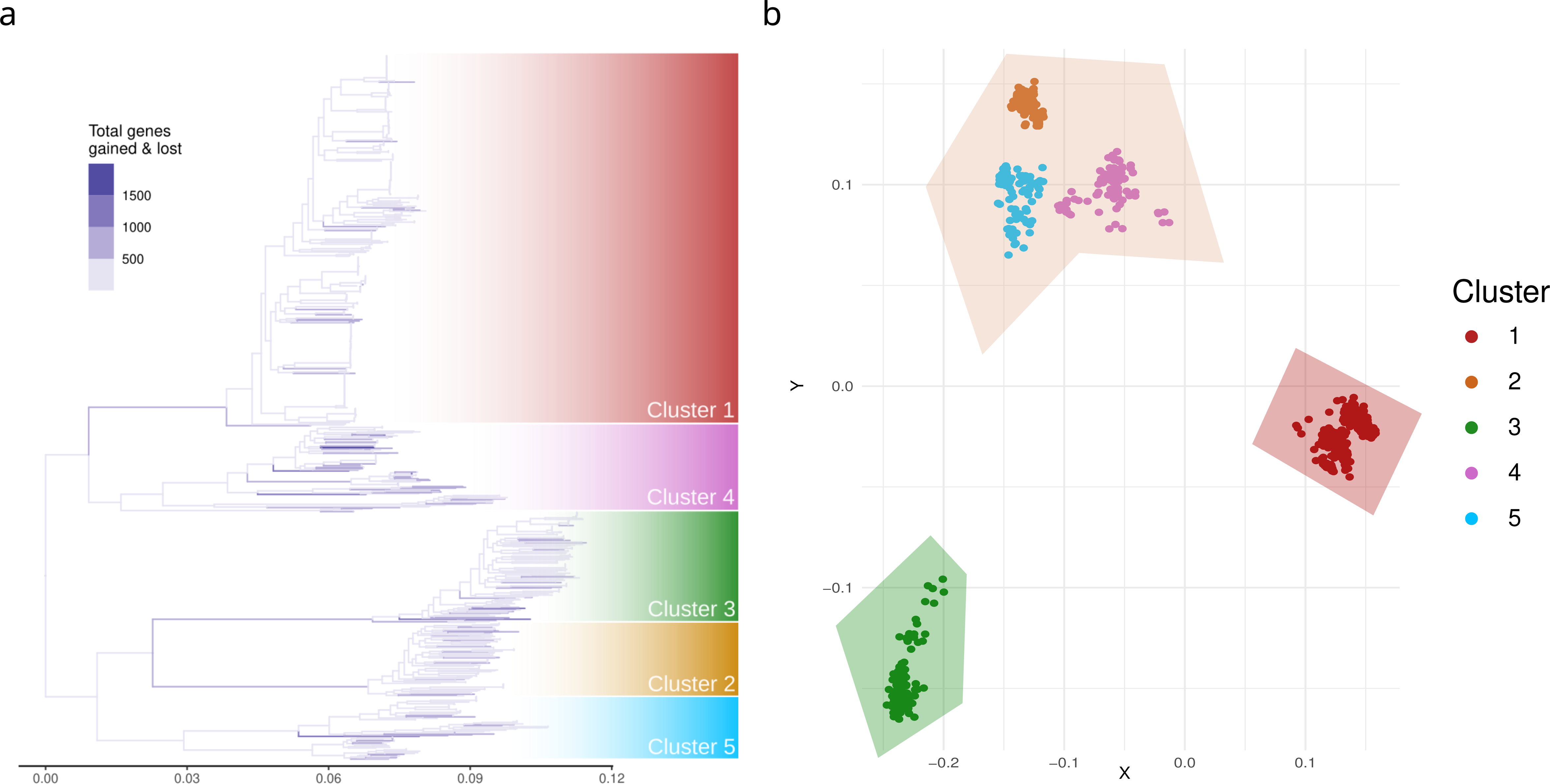
Analysis of the *Serratia marcescens* gene repertoire. a) Phylogenetic tree with branches coloured according to the number of gene gains and losses inferred by Panstripe. b) Principal Coordinate Analysis (PCoA) of *S. marcescens* strains based on gene presence absence. Each dot corresponds to a *S. marcescens* strain coloured on the basis of its cluster. The shaded regions represent the three clusters identified by the K-means unsupervised algorithm.

PCoA on gene presence absence (Figure 3b) clearly groups the strains coherently with the phylogenetic clusters and the K-means unsupervised clustering separates Cluster 1 and Cluster 3, grouping together Clusters 2, 4 and 5 (Figure S8). This result shows that the five *S. marcescens* clusters have distinct gene content, and Cluster 1 and Cluster 3 have, previously found to be associated with clinical and environmental sources, are remarkably different from the others.

A more in-depth analysis identified 107 genes specific for Cluster 1 (i.e. present in >95% of Cluster 1 strains and <15% of the strains of the other clusters), 58 genes for Cluster 2, 168 for Cluster 3, 14 for Cluster 4 and 81 for Cluster 5. Among the genes associated to Cluster 1 (see Table S2) there are *fha*C, with a role in the haemolytic process, *fim*C and *hrp*A, both involved in fimbrial biogenesis and *lac*Z, linked to coexistence with mammals. The genes associated with Cluster 3 included genes involved in the metabolism of plant and fungal carbohydrates (*pul*E, *pul*C, *pul*K and *pul*L). Perhaps, these traits could be linked to the differential ecology of the clusters and offer a genetic hint of their adaptation process. Indeed, *fha*C is essential for pathogenicity in *Bordetella pertussis*^32^ and the acquisition of lacZ, as stated above, can be linked to the association with mammalian hosts^33^. The presence in Cluster 3 of four genes from the *pul* operon could provide a signal of adaptation to plant environments^34^. In summary, at the beginning of their separation, three clusters underwent frequent episodes of gene gain/loss and two of these clusters (Cluster 1 and Cluster 3) reached a unique gene repertoire. Since these clusters were notably found to be enriched in clinical and environmental samples, their gene repertoire is coherent with independent adaptive trajectories towards specific lifestyles. Despite being grouped with Cluster 4 and Cluster 5, also Cluster 2 displays a clear pattern of differentiation in gene content.

### The ecology of *Serratia marcescens* clusters inferred from shotgun metagenomics analysis

As stated above, some clusters present a clear enrichment for specific isolation sources, such as the Cluster 1 for the hospital settings and Cluster 3 for the environment. However, *S. marcescens* is mainly studied for its clinical relevance, producing a strong sampling bias towards hospitals and human samples. To overcome this limit we investigated the presence of strains of the *S. marcescens* clusters in different biomes using a large metagenomics database.

Firstly, we identified protein markers specific to *S. marcescens* and others able to distinguish the clusters. As to *S. marcescens* protein markers, the 40 *S. marcescens*-specific proteins found by Alvaro et al. 2023 (submitted) were tested and 27 resulted to be discriminant. To distinguish the *S. marcescens* clusters, the cluster-specific proteins found above were tested: 46 gene markers were selected for Cluster 1, 11 for Cluster 2 and 20 for Cluster 5. For Cluster 4 and Cluster 3 it was not possible to identify reliable markers. For Cluster 4, the lack of protein markers can be explained by the fact that only 14 cluster-specific core genes were identified (see the “Specific gene repertoires suggest clusters ecological adaptations” section). On the other hand, the absence of specific genes for Cluster 3 can be explained considering the high similarity of its cluster-specific genes with those of other bacterial species (even outside the *Serratia* genus, see Figure S9). This suggests that the peculiar gene content of Cluster 3 could arise from intense gene flow with other bacterial species, coherently with the recently proposed idea that the evolution of the *Serratia* genus is shaped by extensive interspecies gene flow^28^.

To study the distribution of the *S. marcescens* clusters the protein markers were searched into MGnify^35^, a large database containing hundreds of thousands of protein sequences from shotgun metagenomics data on several biomes. The search of *S. marcescens*-specific protein sequences into the Mgnify database identified a total of 6,476 metagenomic-based assemblies possibly containing *Serratia marcescens* sequences. Among these *S. marcescens*-positive assemblies, 5,440 (84%) resulted positive to at least one *S. marcescens* cluster, and 1,720 (32%) resulted positive to a single cluster. Despite a general biome co-presence was observed for the clusters (Figure S10), some interesting statistically significant associations were observed: Figure 4, which only takes most relevant biomes into account, shows that Cluster 1 was enriched in aquatic biomes (i.e. marine and freshwater) and Cluster 2 in the digestive system. (For details on all biomes where *S. marcescens* was traced see Figure S11). This analytical approach presents some issues. Despite the target proteins being selected on the basis of their high specificity, HGT events among *S. marcescens* clusters and between *S. marcescens* and other species cannot be excluded. Indeed, the used protein markers have a sensibility/specificity threshold of 75%. Moreover, metagenomic datasets are highly susceptible to chimeric sequences assembly^36^. Nevertheless, this analysis represents a useful tool to broaden our knowledge on the ecology of *S. marcescens*. It underlines the ecological plasticity in *S. marcescens* and fortifies the idea that, although wide-spread and often co-existent, clusters could have individual ecological preferences. It is of particular interest that Cluster 1 (strongly associated with clinical samples and harbouring virulence factors) was found to be enriched in freshwater, suggesting a possible reservoir for this pathogenic bacterium.

**Figure 4.**
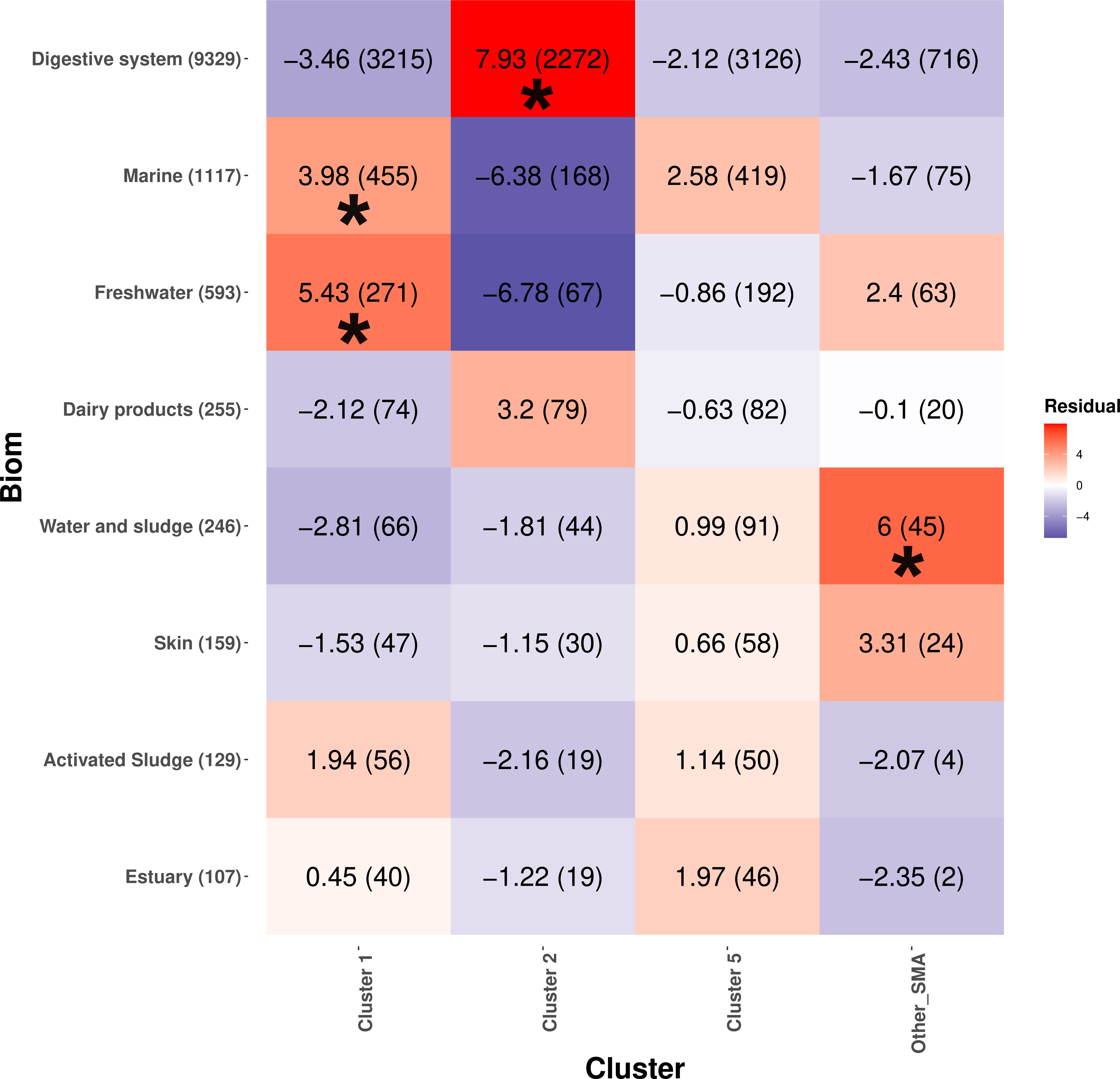
Inference of *S. marcescens* clusters in metagenomic samples. Cluster-specific core genes with high rates of specificity and sensibility (see Methods) were searched on metagenomic samples of the MGnify database to identify phylogenetic clusters of *S. marcescens* in a broad set of biomes. A metagenomic assembly was considered “positive” to a Cluster if >10% of cluster-specific core genes were found within it. When a metagenomic assembly resulted positive to core *S. marcescens* genes, but not to genes associated to Cluster 1, Cluster 2 or Cluster 5, the assembly was defined as “Other SMA”. The heatmap shows the residuals of the Chi-squared test used to investigate whether *S. marcescens* clusters are associated to samples from specific biomes. Statistically significant associations are marked with asterisks (‘∗’, p value <0.05). Here, only biomes with more than 100 positive samples are shown, see Figure S11 for all biomes.

### Reconstruction of the Refined genomic dataset and phylogenetic tree

The analyses used to investigate the *S. marcescens* clusters origin and maintenance (recombination analysis, molecular clock and HGT analysis) are sensitive to genetic dataset biases. For this reason, the Global genomic dataset was refined to balance the *S. marcescens* genetic variability (see Methods and Figure S12). The obtained Refined genomic dataset included a total of 86 representative strains: 19 from Cluster 1, 16 from Cluster 2, 12 from Cluster 3, 21 from Cluster 4 and 18 from Cluster 5. Then, the ML phylogenetic tree was built on the relative 365,317 coreSNPs. The obtained tree was globally coherent with that obtained on the Global genomic dataset and all the clusters corresponded to monophyletic highly-supported groups (bootstrap supports 100, Figure S13).

### Large genomic recombinations contributed to *Serratia marcescens* diversification

The Refined dataset whole-genome alignment and the relative ML phylogenetic tree were subjected to recombination analysis to investigate its impact on the evolution of *S. marcescens*. As a whole*, S. marcescens* exhibited a recombination to mutation ratio (r/m) ratio of 2.35, being significantly less recombinogenic than what estimated for opportunistic pathogens^37^ such as *Salmonella enterica* (r/m = 30.2), *Streptococcus pyogenes* (r/m = 17.2) and *Helicobacter pylori* (r/m = 13.6). Still, this r/m value is comparable to other opportunistic pathogens like *Campylobacter jejuni* and *Haemophilus parasuis,* suggesting that homologous recombination is implicated in the shaping of genetic diversity within *Serratia marcescens*. Furthermore, large recombinations (> 100,000 pb) were mapped on basal nodes of Cluster 2 and Cluster 3, suggesting that the divergence among these two clusters emerged in correspondence of major recombination events. Large recombinations are also evident within Cluster 4 and Cluster 1. Recombination parameters were estimated for each branch of the tree and the distribution of the r/m ratio within the five clusters were compared: Cluster 2 has the highest distribution of r/m ratio and is significantly more recombinant than Cluster 3 (p value < 6.4e-07), Cluster 4 (p value < 4.3e-09) and Cluster 5 (p value < 0.00015). Cluster 1 and Cluster 5 are also significantly more recombinant than Cluster 4 (p value < 0.00693 and p value < 0.01138). Large recombinations along the phylogenetic tree and r/m ratios for each cluster are shown in Figure S14.

Interestingly, the recombinations were not randomly scattered along the genome but there is a 10,000 bp long hyper-recombinated region. This region contains the capsular genes *wza*, *wzb* and *wzc* of the *wz* operon, and a phylogenetic reconstruction of their concatenate has confirmed that they are highly recombined (Figure S15). The bacterial capsule is a well-known virulence factor ^38,39^ and capsular locus have been shown to be recombination hotspots as consequence of immune escaping ^40,41^. This suggests a dynamic interaction with other organisms for all clusters, but could also be linked to the ability to colonise and adapt to diverse ecological niches ^42^.

### Clusters exhibit partial sexual isolation and preferential gene flow routes

Up to here, it was established that the *S. marcescens* population is divided in five well-delimited clusters, emerged also by large recombinations and having specific genetic features, including gene content and recombination rate. To unveil whether preferences in genetic exchange could be involved in the maintenance of genotypic clusters within the species, gene flow within *S. marcescens* was investigated.

The HGT analysis performed on the 1,062 core genes identified 676 events on 443/1,062 (42%) genes. More in detail, 517/676 (76%) HGT events, occurred on a total of 374/443 (84%) genes, involved strains of the same cluster, while 159/676 (24%) HGT events, occurred on a total of 142/443 (32%) genes, involved strains of different clusters (Figure 5a). Among the 517 intra-cluster HGT events, 111/517 (21%) were within Cluster 1, 118/517 (23%) within Cluster 2, 61/517 (12%) within Cluster 3, 113/517 (22%) Cluster 4 and 114/517 (22%) within Cluster 5. The inter-cluster HGT events involved preferentially specific pairs of clusters: most cluster pairs exchanged maximum 1% of the 1,062 core genes, while Cluster 2 - Cluster 4 pair exchanged 68 core genes (> 6%) and Cluster 3 - Cluster 5 pair exchanged 31 genes (> 3%). The preferential trend towards intra-cluster HGT is also evident from the residuals of the Chi-squared test (Figure 5b).

**Figure 5.**
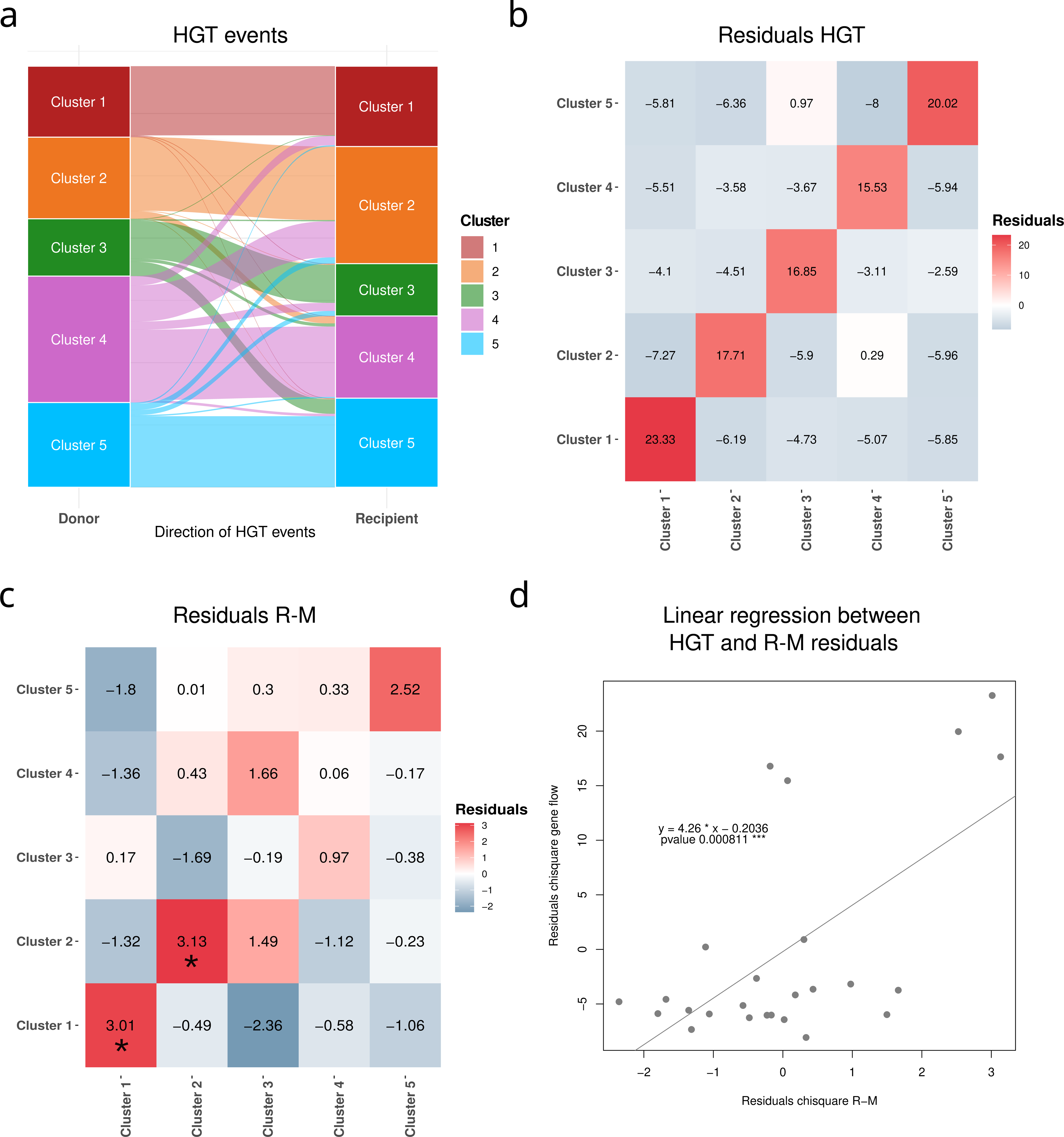
Gene flow and Restriction-Modification (R-M) systems compatibility. a) Stacked bar charts showing the percentage of genes with Horizontal Gene Transfer (HGT) events inferred for each cluster from 1,062 core genes. “Intra” indicates genes with HGT events which happened within strains of the same cluster. “Donor” indicates genes with HGT events departing from the cluster to other clusters. “Recipient” indicates genes with HGT events departing from other clusters. b) Heatmap showing the Chi-squared residuals of *S. marcescens* clusters pairs on the basis of the total number of HGT events inferred between them. c) Heatmap showing the Chi-squared residuals of *S. marcescens* clusters pairs on the basis of R-M systems compatibility. Statistically significant associations are marked with an asterisk (*). d) Linear regression analysis between the residuals of the Chi-squared test on the basis of the R-M compatibility, and the residuals of the Chi-squared test on the basis of the number of HGT events inferred from core genes. Statistically significant associations are marked with asterisks (‘∗’, p value <0.05).

Interestingly, Cluster 1 (associated with clinical samples and a unique gene repertoire) is the most sexually isolated from the other clusters. On the other side, Cluster 4 shows a notable genetic exchange with Cluster 2, despite the phylogenetic distance. Altogether, these analyses revealed a limited gene flow among the clusters, suggesting a sexual isolation coherent with the genetic separation described above.

### Restriction-Modification (R-M) systems could be involved in the genetic isolation of *Serratia marcescens* clusters

One of the main gene flow barriers is the incompatibility between Restriction-Modification (R-M) systems. Bacteria modulate the acquisition of foreign DNA (i.e. avoiding phagic DNA) using two-components Restriction-Modification (R-M) systems: the first enzyme (a methylase) methylates specific DNA patterns, while the second enzyme (an endonuclease) cleaves DNA when the same pattern on the DNA is not methylated. Thus, two bacteria can exchange DNA more successfully if they harbour compatible R-M systems. Genetically close strains tend to harbour similar R-M systems, thus similar strains were grouped and only one representative strain per group was included in the R-M graph (see Methods). A total of 84 strains were selected for R-M graph reconstruction: 30 from Cluster 1, 10 from Cluster 2, 16 from Cluster 3, 20 from Cluster 4 and 8 from Cluster 5. The R-M compatibility among the *S. marcescens* strains of the Global dataset was studied and the R-M compatibility among the clusters was studied by Chi-squared test. The analysis revealed that Cluster 1, Cluster 2 and Cluster 5 have a strong intra-cluster preferential R-M compatibility, as shown by the residuals in Figure 5c. Furthermore, as shown in Figure 5d, the Chi-square Test residuals obtained by the analysis of R-M systems resulted significantly correlated to those obtained by HGT analysis (Linear regression, p = 0.0008). This result suggests that the observed sexual isolation of the clusters could be partially explained by genetic barriers due to R-M system incompatibility.

## Discussion

*Serratia marcescens* is an infamous nosocomial pathogen able to cause large and fatal outbreaks in Neonatal Intensive Care Units (NICUs) and to rapidly spread in hospital settings^43,44^. The bacterium is also able to colonise soil, water, plants and animals such as insects and corals. Recent genomic studies have identified clades associated with different isolation sources, including clinical/hospital-based lineages harbouring several virulence/resistance traits^24–28^.

In this work, we investigated the diversity of this bacterium, with a strong focus on the genetic/ecological mechanisms underlying its population structure. Our results clearly showed the existence of five well-defined clusters that exhibited specific gene content and limited inter-cluster gene flow. At least two clusters also showed genetic signatures of ecological adaptations. Cluster 1 is frequently associated with hospital settings: it has one of the most reduced pan-genome sizes, high rates of gene gain/loss in its phylogenetic basal node and it comprises a very specific gene repertoire, including genes involved in virulence mechanisms. This cluster also has reduced intra-cluster genomic variability and a very limited gene flow with other clusters. Overall, these genetic features are compatible with an association to a hospital-related lifestyle. Interestingly, metagenomics analysis revealed that Cluster 1 could use water (marine or freshwater) as its reservoir habitat. This environmental association suggests a possible explanation of the ability of strains belonging to this cluster to rapidly spread in the hospital environment and to colonise several substrates, but further studies are required to test this hypothesis. Cluster 2 also displays several events of gene gain and loss at its root, which produced a gene composition distinct but not so distant from Cluster 5 and Cluster 4. It is not associated with any isolation source, but the metagenomic analysis suggests its enhancement in the human digestive system. Cluster 3 is enriched in environmental sources (such as soil, plants and water) and displays a specific gene repertoire which includes genes involved in the metabolism of plant and fungal carbohydrates. The cluster has also a reduced genetic variability and it displays signals of an extensive gene flow with other bacterial species. Cluster 4 is not associated with any isolation source. It is characterised by large genetic variability, greater genome size and the widest pan-genome, with very few specific genes. Cluster 5 is enriched in environmental and animal sources but the genetic variability within the cluster is high and gene composition is very similar to Cluster 4. Moreover, the metagenomic analysis did not reveal a specific association for any biome. Interestingly, it is the only cluster with a macro geographic uneven distribution (association to North America).

The existence of deeply demarcated clusters in *S. marcescens* suggests geographical, ecological or genetic barriers behind the origin and maintenance of this diversity. The analysis of isolation sites strongly supports the absence of geographical barriers: strains of all the clusters have been often isolated in the same hospital in the same period of time, in our dataset as well as in the study published by Moradigaravand and colleagues^45^.

Considering this limited geographical isolation, the potential ecological divergence between the clusters was investigated using genomic information. More in detail, gene content of each cluster was compared to trace signals of ecological adaptation^8^ and, when available, specific genetic traits were searched in a large metagenomic database to better understand cluster ecology and evade sampling biases. Interestingly, Cluster 1 and Cluster 3 showed consistent signals coherent with independent adaptive trajectories to hospital and environmental settings, respectively. This result partially recalls the idea first proposed by Abreo and Altier in 2019^24^, that *S. marcescens* has diverged into an “environmental” clade and a “clinical” clade. However, unlike suggested by Abreo and Altier, the divergent “ecotypes” represent two emerging clades and only represent a minimal portion of the genetic variability observed in *S. marcescens*.

The absence of strongly evident ecological adaptation for each of the clusters led us to investigate the origins of these clusters in light of recombinations and gene flow. The clusters mainly show an average recombination rate, in comparison to other species, even if several large recombination events were detected, within the clusters but also on the basal nodes of sister-groups Cluster 2 and 3. The reconstruction of horizontal gene transfer (HGT) events on core genes revealed a very limited inter-cluster gene flow, suggesting the presence of ancient and strong barriers to recombination. Furthermore, the analysis of Restriction-Modification (R-M) systems revealed a partial incompatibility among clusters which could eventually explain the observed genetic isolation. All these results suggest that genetic barriers have had a major role in the sexual isolation of the clusters, shaping the current population structure of *S. marcescens*.

In conclusion, *S. marcescens* is composed of sexually-isolated clusters separated by strong genetic barriers. Within this population structure, two clusters (Cluster 1 and 3) have initiated adaptive trajectories to specific ecological niches and proceed to progressively isolate from the others. Whereas, other clusters are ecologically generalist and despite they often co-occur in the same environment at the same time, genetic barriers are sufficiently thick to maintain the clusters regardless of ecology or spatial distribution. Thus, with a hint of speculation, we propose that the leading role in the evolution of *S. marcescens* is played by the genetic barriers between co-occurring, ecologically generalist subpopulations. Eventually, due to environmental pressure and constant reshuffling of the accessory genome with other species, adaptive populations have emerged. Our results open to interesting biological questions, such as: what caused the arisal of genetic barriers in the first place? Is this cluster-like population structure in equilibrium or are the adaptive clusters embarked on paths towards speciation? At what point could these clusters be considered as subspecies?

## Methods

### Global genomic dataset preparation

#### Preliminary genome dataset reconstruction

The preliminary genomic dataset used in the study contained a total of 1,113 genome assemblies: i) 871 *Serratia marcescens* genomes available on March 10, 2022 in the Bacterial and Viral Bioinformatics Resource Center (BV-BRC) for which geographical information and isolation date were reported; ii) seven additional genomes used in previous genomic studies^24,25,45^ and absent in the BV-BRC database; iii) 230 *S*. *marcescens* genome assemblies sequenced in the previous study (Alvaro et al. 2023, submitted); iv) five *S. marcescens* genome assemblies of strains isolated from the Italian hospitals ASST Papa Giovanni XXIII Hospital in Bergamo (n=2), RCCS San Raffaele Hospital (HSR) in Milan (n=2) and ASST Fatebenefratelli Sacco Hospital in Milan (n=1) (details about genome sequencing and assembly are reported below). Details are reported in Table S1.

#### Serratia marcescens genomes sequencing and assembly

Five *S. marcescens* isolates were grown on McConkey agar medium overnight at 37°C. The day after, single colonies were picked and DNA extractions were carried out using a Qiagen QIAcube Connect automated extractor (Qiagen, Hilden, Germany) following the “bacterial pellet” protocol which employs Qiagen DNeasy Blood & Tissue reagents. Then, libraries were prepared and 2 × 150 bp paired-end run sequencing was carried out on the Illumina NextSeq platform. The reads were quality checked by using FastQC tool (https://www.bioinformatics.babraham.ac.uk/projects/fastqc/) and then assembled using SPAdes ^46^.

#### Global genomic dataset selection

Within the preliminary genome dataset, the low quality genome assemblies and those for which the *S. marcescens* taxonomy was incorrect were detected and removed to obtain the “Global genomic” dataset.

The assembly quality parameters used for the selection were: assembly total length, number of contigs, N50, N count and the Open Reading Frame (ORF) number. ORF calling was performed using Prodigal ^47^ and the genome statistics were obtained using the assembly-stats tool (https://github.com/sanger-pathogens/assembly-stats). For each of these parameters, the thresholds for the selection were computed on the starting genomic dataset using the Tukey’s fences statistical method^48^: the lower boundary (L) is computed as ▭1 − (1.5 ▭ ▭▭▭) and the higher boundary (H) as ▭3 + (1.5 ▭ ▭▭▭), where Q1 indicates the first quantile of the value distribution, Q3 indicates the third quartile and IQR indicates the interquartile range. The obtained thresholds were: i) “Total length” between 4,500,000 bp and 6,000,000 bp; ii) “Number of contigs” < 116; iii) “N count” < 5,842 ; iv) “N50” > 7,077 ; v) 5,134 < “ORF count” < 4,594. The “N count” parameter was considered crucial for high-quality and all genomes that did not respect its threshold were excluded. Among the remaining genome assemblies, those that passed at least three out of the other four quality checks were selected for the taxonomy-based step of selection.

Taxonomy of the genomes were assessed combining Average Nucleotide Identity (ANI) and 16S rRNA sequence. The Mash pairwise distance matrix was computed between all genomes using Mash^29^ and the genomes were clustered with a cut--off distance of 0.05. The 16S rRNA sequence was extracted using Barrnap and Blastn-searched into the 16S rRNA database Silva^49^: the genomes were then classified on the basis of the best hit as “*Serratia marcescens*”, “*Serratia* spp.” and “Others”. The 16S rRNA gene is in multiple copies within the *S. marcescens* genome making it difficult to assemble. Genomes for which it was not possible to identify the 16S rRNA gene were classified as “undefined”. Combining the Mash-based clustering and the 16S rRNA classification, a *Serratia marcescens*-like cluster was defined. The genomes clustered within the *Serratia marcescens*-like cluster and annotated with 16S rRNA as “*Serratia marcescens*”, “*Serratia* spp.” or “undefined” were selected. Herein, the selected genome dataset will be referred to as the “Global genomic dataset”.

#### Genome classification by origin

Based on the sampling material, *S. marcescens* genomes were manually distinguished into three ecological categories: i) *clinical*, if the sample was obtained from a clinically--related human sample; ii) *animal*, if the bacterium was associated to any non--human metazoan; iii) *environmental*, if the sample was found on any other environmental source, such as water, plants and soil.

### Population structure

#### SNP-based phylogenetic analysis

The assemblies of the Global genomic dataset and one outgroup (*Serratia plymuthica strain 4Rx13,* GCF_000176835.2) were aligned against the genome assembly of the reference strain *S. marcescens* Db11 and Core Single Nucleotide Polymorphisms (CoreSNPs) were called using the tool Purple^50^. The obtained CoreSNPs were subjected to Maximum Likelihood (ML) phylogenetic analysis using FastTree MP^51^ (with 100 pseudo-bootstraps), using the general time reversible (GTR) model. The obtained tree was manually rooted on the outgroup using Seaview^52^. Lastly, the web--based tool iTOL^53^ was used to map strains metadata on the topology.

#### Clustering analysis

The global genomic dataset *strains were* grouped via Principal Coordinates Analysis (PCoA) and unsupervised clustering algorithm K-means, using independently tree patristic distances, CoreSNP distances, Mash distances and Jaccard distances computed on the gene presence absence. The Average Nucleotide Identity (ANI) between strains was computed as (1 - Mash distances) * 100 ^29^. For each analysis, the optimal number of clusters was determined in accordance to the best average silhouette score.

### Clusters comparison

#### Genomic features

Genome size, number of genes and GC content were compared between clusters by Mann-Whitney U-test with Holm post-hoc correction and visualised by boxplots. Pairwise SNP-distances and ANI distances were used to infer genetic diversity among strains and compared among the clusters by histograms. The analyses were performed using R.

#### Synteny

The genomic synteny within and between clusters was assessed on the complete genomes available in the global genomic dataset. Before the analysis, plasmidic contigs were manually removed and the chromosomes were re-arranged on the basis of the *dnaA* gene position. For each cluster the re-oriented genome assemblies were aligned using progressiveMauve^54^ and the intra-cluster synteny plot was obtained using the R package “genoplotR”^55^. The inter-cluster synteny was investigated using one representative strain per cluster.

#### Cluster geographic distribution

The geographic distribution of the strains of the different clusters was compared using the Chi-squared Test of Independence on the isolation continents. Pearson’s standard residuals were evaluated to investigate geographic distribution of the clusters: i.e. residuals were considered as statistically significant when the value was greater than the Bonferroni-corrected critical value ^56^.

#### Cluster sampling source enrichment

As stated above (i.e. section “Genome classification by origin”), the *S. marcescens* strains of the Global genomic dataset were assigned to ecological categories on the basis of their isolation source. To investigate ecological preferences among clusters, the Pearson’s standard residuals of the Chi-squared Test of Independence between *S. marcescens* cluster and the relative ecological categories were studied. The residuals were considered as significant if their absolute value was greater than the Bonferroni-corrected critical value. To minimise the possible bias due to geographical proximity of the samples, a geographically-balanced Chi-squared Test of Independence was implemented with a Monte Carlo method: the test was run 1,000 times, sampling 40 genomes from each continent. A cluster was considered statistically associated to a specific ecological category when the relative standard residual was significant in at least 950 test runs out of 1,000 replicates. Strains from Africa (n = 8) and Asia (n = 3) were excluded from the analysis because of the very low representation of these continents in the dataset; North America (n = 96) and South America (n = 22) were merged into the “America”.

#### Pan-genome analysis

Genomes were annotated using Prokka^57^ and General Feature Format (GFF) files were fed to Roary^58^ for pan--genome analysis. Pan-genome cumulative curves were built using R for the entire dataset and for each cluster independently. Then the open vs close status of each pan genome was assessed as described by Tettelin et al.^59^ Gene gain and loss events were mapped on the tree with Panstripe^60^, using maximum parsimony as method for the ancestral state reconstruction.

Differences in gene content among clusters were also investigated by PCoA on the gene presence/absence Jaccard distance matrix obtained from the Roary tool.

#### Annotation of cluster-specific core genes

Orthology groups that were found to be core (> 95% present) in one cluster and rare in all other clusters (<15% present) were considered cluster-specific core genes. Nucleotide distances among sequences of each orthology group were computed via the *dist.dna* function of the “ape” R library^61^, and the sequence with the lowest mean nucleotide distance from the others was selected as representative of each orthology group. Representative sequences were annotated against the COG-database using the tool COGclassifier (https://pypi.org/project/cogclassifier/). Moreover, genes were defined as chromosomal or plasmidic by BLAST search against the complete genomes with plasmids included in the dataset.

#### Investigation of the S. marcescens clusters enrichment in ecological niches using a large shotgun metagenomics database

The analysis of ecological enrichment performed on the Global genomic dataset (see above) can suffer from sampling bias. Indeed, as expected, most of the strains were isolated from clinical settings. The Mgnify^35^ protein database contains protein sequences obtained from shogun metagenomics sequencing of thousands of samples collected from a vast range of biomes/ecological sources. To assess the presence of sequences specific to the different *S. marcescens* clusters into the samples this database can help to overcome this issue.

To do so, it was necessary to use protein markers able to discriminate the *S. marcescens* clusters from all the other bacterial species. The protein sequences of the cluster-specific core genes (see section “*Annotation of cluster-specific core genes*”) were searched by DIAMOND^62^ (E-value < 0.00001, sequence identity >= 90% and the ratio between query length and length of the hit >= 0.85 <= 1.1) against all proteins of the genomes of the Global genomic dataset, in order to assess the ability of these target proteins to identify the clusters by DIAMOND search. For each target protein, the sensibility and specificity was evaluated using the Youden’s index and the best threshold for the percentage of sequence identity was determined in a range from 90 to 99. The specificity of the target proteins for *Serratia marcescens* was assessed in a similar way: protein sequences were searched by DIAMOND against the NCBI NR^63^ database. After filtering for coverage and e-value as above, hits were categorised on their taxonomy as hits on *S. marcescens* or as and the same set of percentage of sequence identity values was used. The highest value of specificity was extracted together with the corresponding percentage of sequence identity used as threshold. The protein sequences of 40 core genes with a good specificity for *S. marcescens* determined by Alvaro *et al.* 2023 (submitted) were also included in this analysis. The genes were selected to be appropriate markers only if, at a certain threshold, their Youden’s index value was higher than 0.75 and the specificity to *S. marcescens* higher than 75%.

All sequences of the marker genes were searched by DIAMOND against the Mgnify protein database. The results were filtered as above for coverage and e-value, while the percentage of sequence identity threshold used was target-specific. A Mgnify sample was considered to contain *S. marcescens* if at least three of the selected Alvaro *et al.* 2023 (submitted) protein targets were present. These samples were further investigated for the determination of the *S. marcescens* cluster present by DIAMOND searching for the cluster-specific protein markers. To determine if *S. marcescens* clusters were linked to different biomes/ecological sources, Chi-Squared Test of Independence was performed (standard residuals were considered significant if their absolute value was greater than the Bonferroni-corrected critical value).

### Cluster gene flow and recombination analysis

#### Dataset refinement

The analyses for the investigation of cluster origin and maintenance (including recombination, gene flow and molecular clock analyses) are sensitive to the size and genetic bias of the genomic dataset. To reduce the size of the Global genomic dataset, maintaining the genetic variability as much as possible, the genomes were grouped on the basis of pairwise coreSNP distance: the strains having coreSNPs distance below a specific threshold fell in the same group and the youngest and oldest (on the basis of the isolation date) strains of the group were retrieved. To define the best threshold to be used, the number of groups over SNPs thresholds ranging between 0-1,000 SNPs were plotted using R. Herein, this dataset will be referred to as “Refined genomic dataset”.

#### SNP calling, SNP annotation and Maximum Likelihood phylogenetic analysis

The Purple^50^ tool was used for the reference-based coreSNP calling. The genome of the Refined genomic dataset was aligned to the *S. marcescens* Db11 reference genome assembly and SNP were called and used to obtain the whole-genomes alignment and to extract the coreSNPs^50^. The extracted coreSNPs were then subjected to Maximum Likelihood (ML) phylogenetic analysis using RAxML8^64^, applying a general time reversible model that incorporates rate variation among sites and proportion of invariant sites (GTR + G + I), according to ModelTest-NG^65^.

#### Whole-genome recombination analysis

The ML phylogenetic tree and the whole-genome alignment (obtained using Purple^50^) were fed to ClonalframeML^66^ for recombination analysis. Recombination events were estimated per-branch and ambiguous sites on the alignment were ignored in the analysis. From ClonalframeML output, r/m ratio was calculated as ▭/▭ = ▭/▭▭▭▭▭ * ▭▭▭▭▭ * ▭▭ and compared among clusters via Mann-Whitney U-test with Holm post-hoc correction. Then, for each cluster the cumulative number of recombined bases within windows of 5-kbp along the whole-genome alignment was computed.

The gene annotation of reference genome Db11 was checked to identify genes located on highly recombined regions. The ML phylogeny of the genes of interest within the recombined regions were obtained using RAxML8 with 100 pseudo-bootstraps after best model selection using ModelTest-NG. Moreover, single large recombination events along the genome (> 100 kbp) were mapped on the phylogenetic tree.

#### Gene flow analysis

Core gene alignments were extracted from the whole-genomes alignment obtained above (see SNP calling, SNP annotation and Maximum Likelihood phylogenetic analysis) on the basis of the positions of the Coding DNA Sequences (CDSs) on the reference Db11 *S. marcescens* strain genome. Each gene alignment was subjected to ML phylogenetic analysis using RAxML8 after best model selection using ModelTest-NG. The topology of each tree was compared to the SNP-based phylogenetic tree using T-REX command-line version^67^ to detect HGT: the analysis was repeated on bootstrap trees and only HGT events with a bootstrap support of at least 75 were considered reliable. The HGT analysis returns the nodes of donors and recipients of each detected HGT event. Using this information, the network describing the gene flow between clusters in *Serratia marcescens* was constructed using Gephi^68^. Lastly, the preferential association among clusters for HGT events was evaluated analysing the residuals of the Chi-Squared Test.

#### Analysis of Restriction-Modification (R-M) systems compatibility among the clusters

The methylase and endonuclease enzymes of the R-M systems present in the strains of the Global genomic dataset were identified and annotated by Blastn search against the REBASE database^69^, selecting the best hits with coverage (hit length / query length) > 0.9 and nucleotide identity > 90%. The hits were then classified as “Orphan methyltransferase” (methylases without the relative endonuclease, usually involved in gene regulation), “Methyltransferase” and “Endonuclease”. When the HGT donor strain harbours methylase enzymes compatible with the endonuclease enzymes of the recipient strain (i.e. the two enzymes methylate/not-cleave the same DNA pattern), the transferred DNA is more likely to be incorporated by the recipient. Strains genetically very similar will tend to harbour similar R-M systems, because they share a closer ancestor and it is reasonable that this could lead to the overestimation of the intra-cluster R-M compatibility. To avoid this bias the strains were previously clustered using the coreSNP alignment obtained above with a threshold of 10 SNPs. Among the strains of the same cluster the one harbouring more R-M system genes was selected as representative. These selected strains were then used to reconstruct the R-M graph where the nodes are the strains, and two nodes are connected if all the R-M systems harboured by the strains are compatible. The preferential association of the clusters on the graph was studied by Chi-squared Test of Independence.

To investigate whether the R-M system could affect the observed gene flow pattern (as determined above), the Chi-square Test residuals of the preferential association between clusters computed from the R-M graph and the residuals obtained from HGT analysis were compared by linear regression.

## Supplementary captions

**Figure S1. K-means clustering on core Single Nucleotide Polymorphisms distances.**

a), c) and e) Silhouette plot of the number of clusters on the basis of core Single Nucleotide Polymorphisms (coreSNP) distances, Mash distances and patristic distances between *S. marcescens* strains. The number of clusters with the highest score was chosen as the optimal number of clusters to fit the data in. b), d) and f) Principal Coordinate Analysis (PCoA) performed on SNP distances, Mash distances and patristic distances, coloured on the basis of the clusters identified by the K-means algorithm.

**Figure S2. GC content and genome length in *S. marcescens* clusters.**

a) Boxplot showing the percentage of GC content in the genomes of each cluster. The p-value of the Kruskal-Wallis test, performed to test the variance between groups, is shown on the bottom left. The groups with significant pairwise differences (Mann-Whitney U test) are connected and the p-value is written on top.

b) Boxplot showing the genome sizes of each cluster. Kruskal-Wallis test and Mann-Whitneu U test are reported as above.

**Figure S3. Synteny between *S.marcescens* clusters.**

Visualisation of the synteny between the 65 complete genomes present in the study. The synteny is shown for strains within each cluster. On the bottom right (“All clusters”), the synteny between clusters is visualised using one genome representative of the most common syntenic variant for each cluster.

**Figure S4. Macrogeographical distribution of *S.marcescens* clusters.**

a) Pie charts representing the frequency of each cluster in the six continents (starting from the left: on top North America, Europe, Asia; at the bottom South America, Africa, Oceania). The total number of strains isolated in the continent (n) is shown under each pie chart. b) Chi-squared test was used to assess whether *S. marcescens* clusters are associated with geographical continents. The heatmap shows the Pearson residuals of the Chi-squared test and statistically significant associations are marked with an asterisk (*).

**Figure S5. Spatiotemporal distribution of *S. marcescens* in Italian hospitals.**

Plot showing the spatiotemporal distribution of 235 *S.marcescens* strains sampled from six Italian hospitals.

**Figure S6. Pan-genome curves for *S.marcescens* clusters.**

Pan-genome accumulation curves for each cluster of *S. marcescens* strains. The region between 1 and 300 strains is zoomed to allow a better visualisation of curves belonging to smaller clusters. Clusters are coloured according to the key on the right.

**Figure S7. Analysis of the pan-genome openness for *S.marcescens* clusters.**

The plot shows the log-log regression for new genes found in an increasing number of genome sequences within *S.marcescens* clusters. A slope of the regression line < 1 indicates that when more genomes are analysed, the number of new genes decays slowly, thus the pan-genome is open.

**Figure S8. K-means clustering on gene presence absence-based Jaccard distances.**

a) Silhouette plot of the number of clusters on the basis of Jaccard distances computed from gene presence absence between *S. marcescens* strains. The number of clusters with the highest score was chosen as the optimal number of clusters to fit the data in. b) Principal Coordinate Analysis (PCoA) performed on Jaccard distances computed from gene presence absence, coloured on the basis of the clusters identified by the K-means algorithm.

**Figure S9. Sequence similarity between proteins of cluster-specific core genes and sequences outside *S. marcescens*.**

Boxplot showing the percentages of nucleotide identity obtained comparing protein sequences of cluster-specific core genes against protein sequences not belonging to *Serratia marcescens* on the NCBI nr database.

**Figure S10. Presence of *S. marcescens* clusters in metagenomic samples from different ecological sources/biomes**

The presence of Cluster 1, Cluster 2, Cluster 5 and other S. *marcescens* was investigated in metagenomic samples of the MGnify database searching for cluster-specific core genes and *S. marcescens* core genes. The heatmap shows the frequency with which every combination of clusters was found in samples from different ecological sources/biomes. In brackets, the number of samples for each combination.

**Figure S11. Inference of *S. marcescens* clusters in metagenomic samples**

a) The stacked bar chart shows the biomes of metagenomic assemblies from the MGnify database in which *S. marcescens* clusters were identified. An assembly was considered positive to a Cluster if >10% of cluster-specific core genes were found within it. When a metagenomic assembly resulted positive to core *S. marcescens* genes, but not to genes associated to Cluster 1, Cluster 2 and Cluster 5, the assembly was defined as “Other SMA”. b) The heatmap shows the residuals of the Chi-squared test used to investigate whether *S. marcescens* clusters are associated to samples from specific biomes. Statistically significant associations are marked with asterisks (‘∗’, p value <0.05).

**Figure S12. Determination of the optimal coreSNP threshold to create the Refined genomic dataset**

In order to reduce the size of the dataset and the genetic bias (for recombination, gene flow and molecular clock analyses) the 902 *S. marcescens* genomes were grouped in coreSNP groups on the basis of a threshold in pairwise coreSNP distance. The determination of the threshold was optimised to reduce and balance the dataset as much as possible with a minor impact on the represented genetic variability. a) To evaluate the potential loss of genetic variability, the mean coreSNP distance for each cluster between strains was plotted for every coreSNP threshold between 0 and 1000 b) To evaluate the size reduction of the dataset, the number of coreSNP groups was plotted for every coreSNP threshold between 0 and 1000. On the basis of both plots, a threshold of 500 coreSNPs was considered optimal.

**Figure S13. Unrooted phylogenetic tree of *S. marcescens* strains in the Refined genomic dataset**

SNP-based Maximum Likelihood (ML) phylogenetic tree (performed with 100 pseudo-bootstrap) including the 86 *S. marcescens* strains of the Refined genomic dataset. The five clusters correspond to five monophyletic groups. Bootstraps above 90 are represented as black dots on the corresponding node.

**Figure S14. Large recombinations and recombination to mutation ratio in *S. marcescens* clusters**

a) Plot illustrating large recombination events (>50 kbp) along the genome (on the right) mapped on every node of the phylogenetic tree (on the left). The colours on the phylogenetic tree represent the *S. marcescens* clusters and the dashed red lines indicate the basal node of each cluster. For Cluster 4, the basal node does not correspond to any large recombination. b) Boxplot showing the recombination to mutation ratio (r/m) for the nodes of each cluster. The p-value of the Kruskal-Wallis test, performed to test the variance between groups, is shown on the bottom left. The groups with significant pairwise differences (Mann-Whitney U test) are connected and the corresponding p-value is written on top.

**Figure S15. Recombinations along the *Serratia marcescens* genome in each cluster**

On the left, distribution of cumulative recombination events along the *Serratia marcescens* genome. The analysis has been performed on a multi-genomes alignment of 86 strains selected to be representative of the clusters (genome alignment was obtained using the genome of *S. marcescens* Db11 strain as reference, position on the x-axis refer to this genome assembly). For each cluster, the recombination distribution is shown as the cumulative number of recombinated bases for genomic windows of 5 kbp. A 10-kbp long highly-recombinated region, containing the capsular genes *wza*, *wzb* and *wzc*, is highlighted by the red box. On the right, comparison among the Maximum Likelihood (ML) phylogenetic trees of the 86 strains, obtained using coreSNPs (on the left) and *wza-wzb-wzc* concatenate (on the right). The results highlight that capsular genes are recombination hotspots.

