## Supplementary figures and images for "Genetic barriers more than ecological adaptations shaped *Serratia marcescens* diversity"

### Figure S1

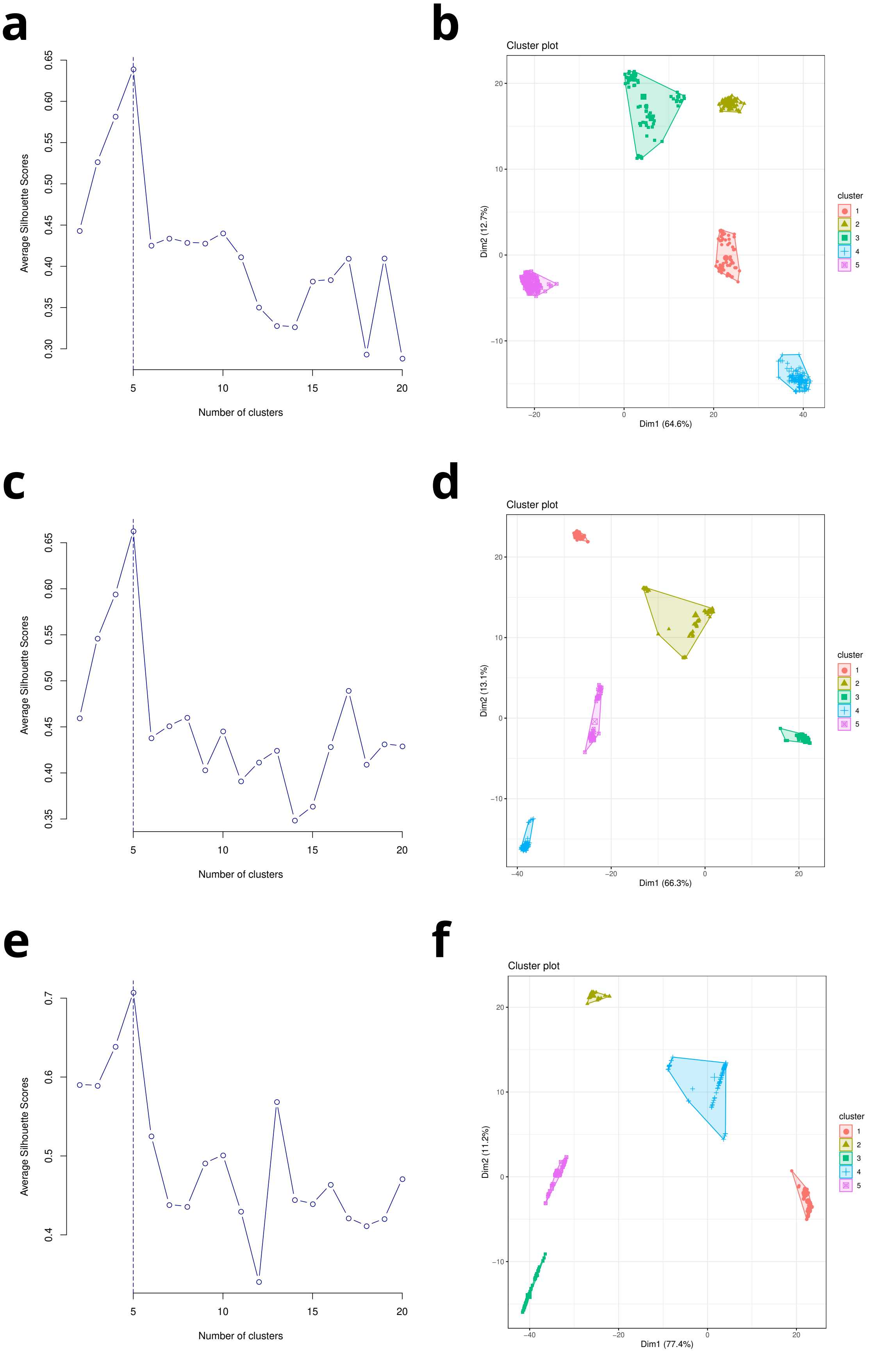

### Figure S2

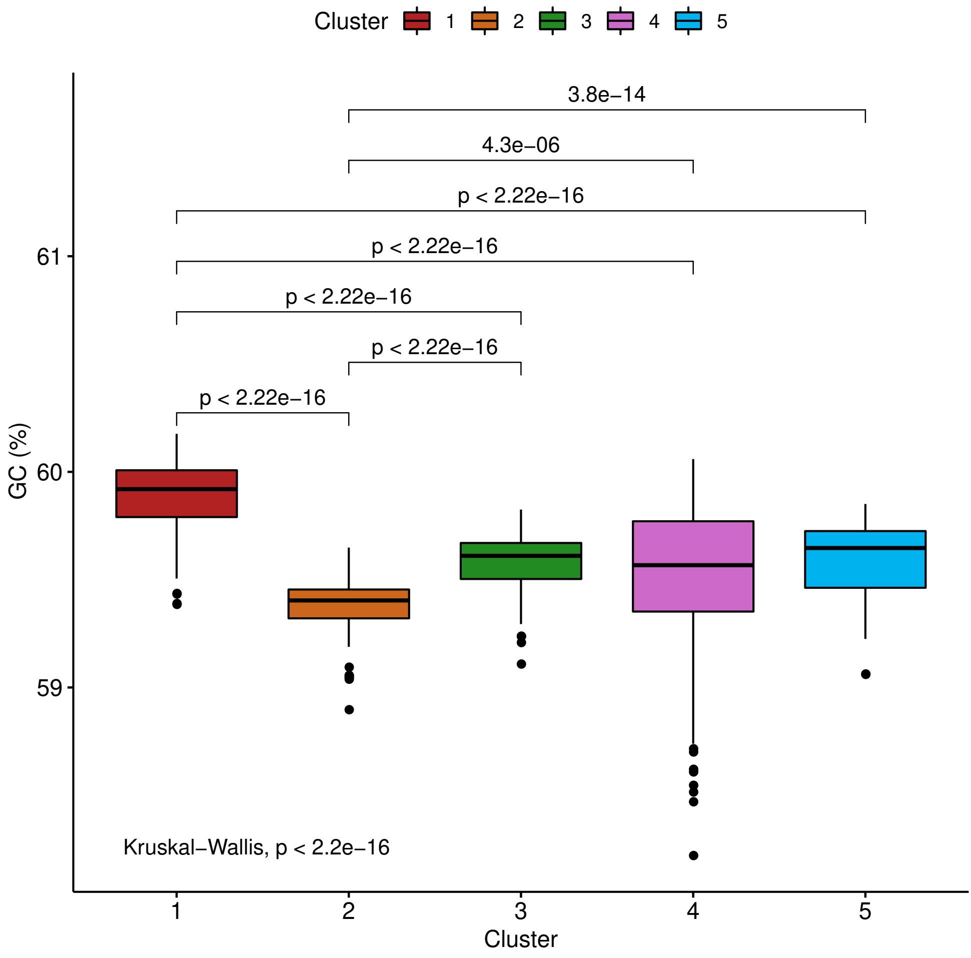

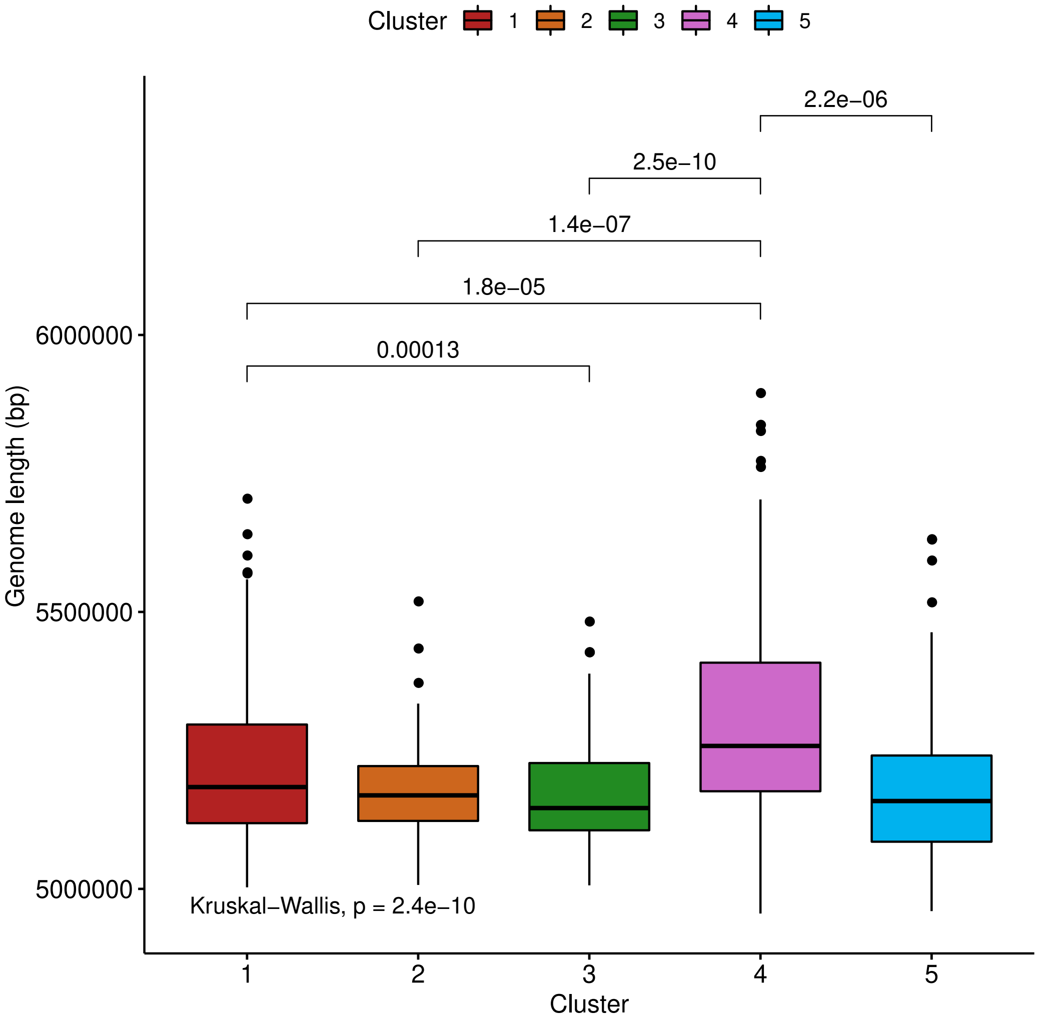

### Figure S3

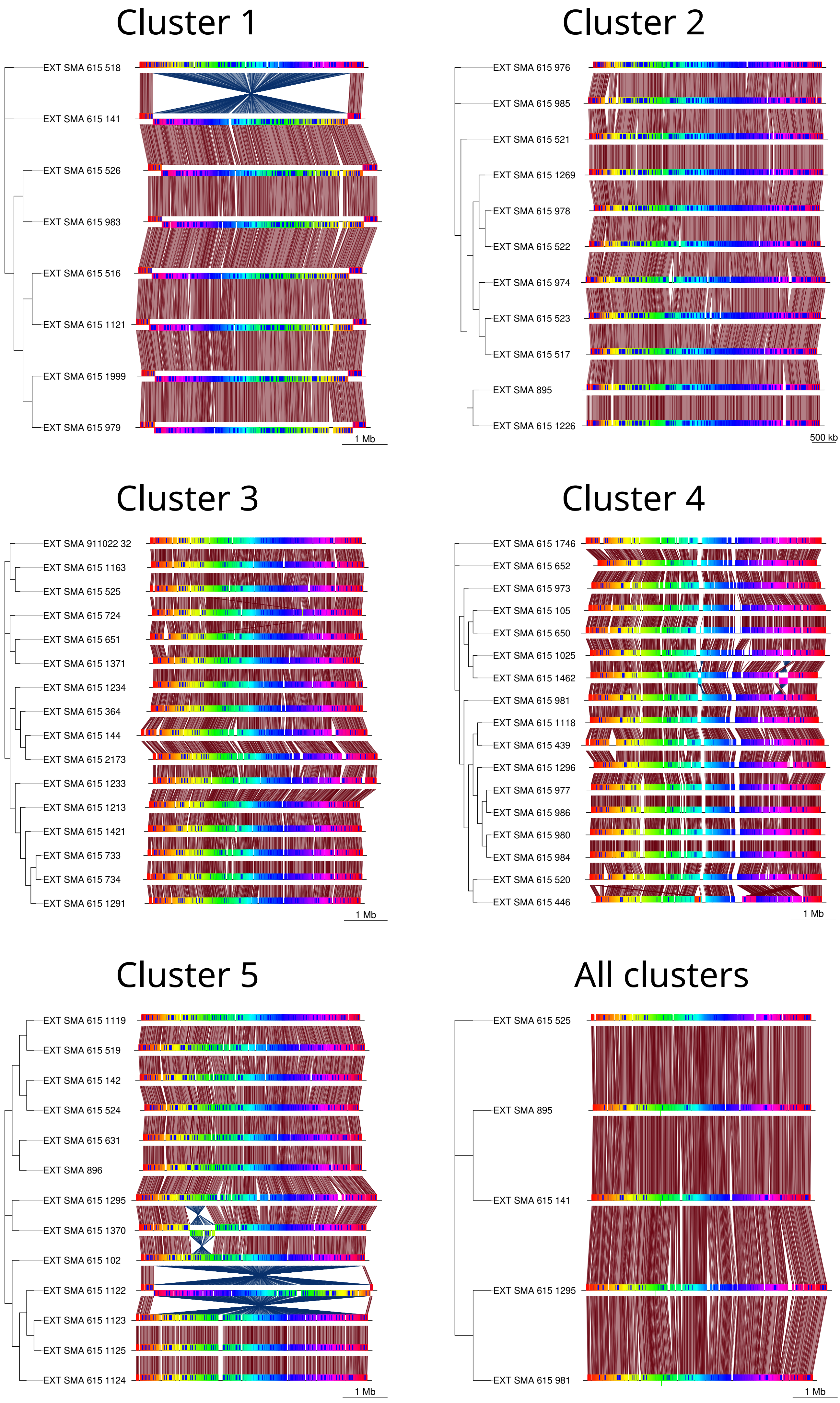

### Figure S4

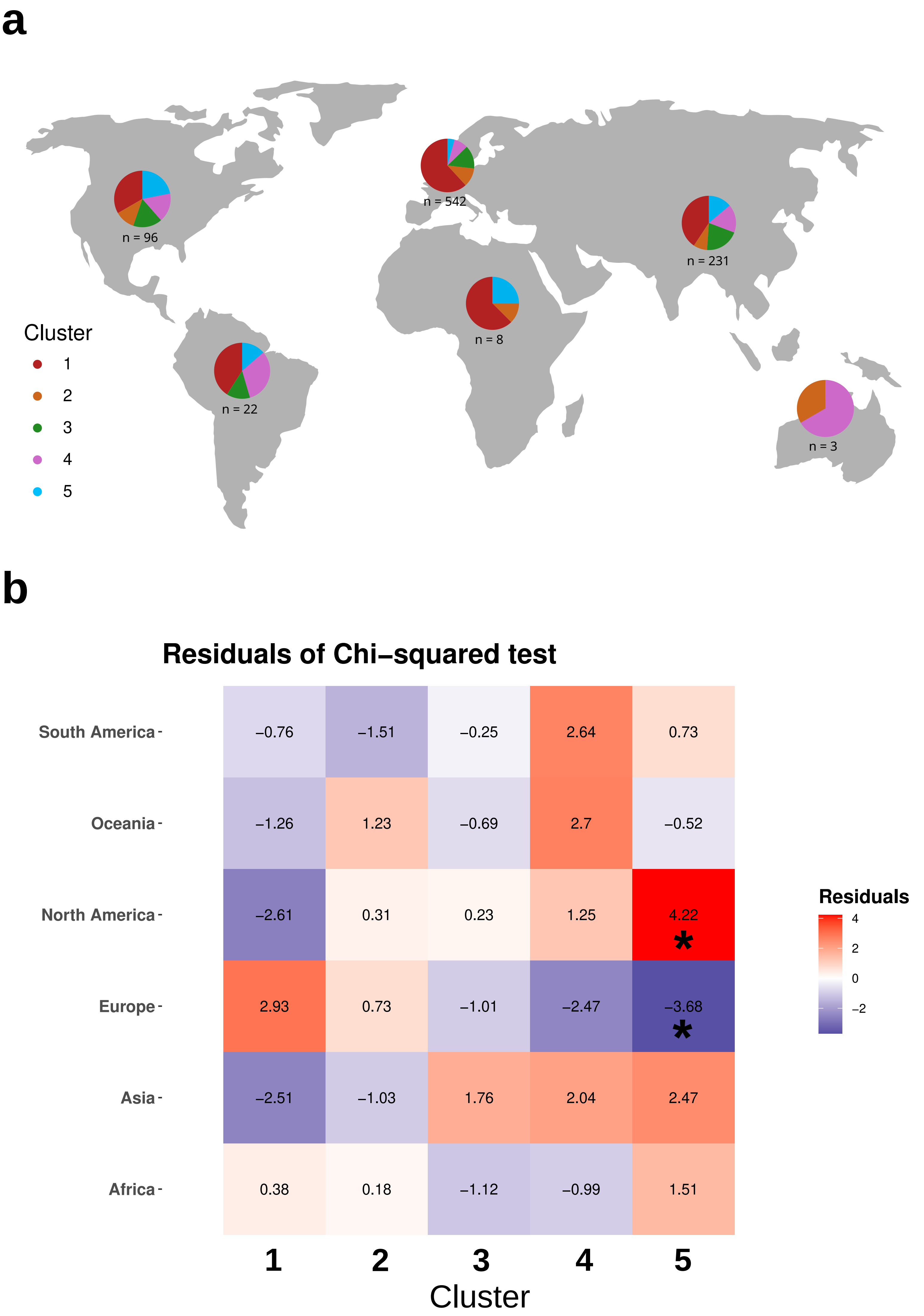

### Figure S5

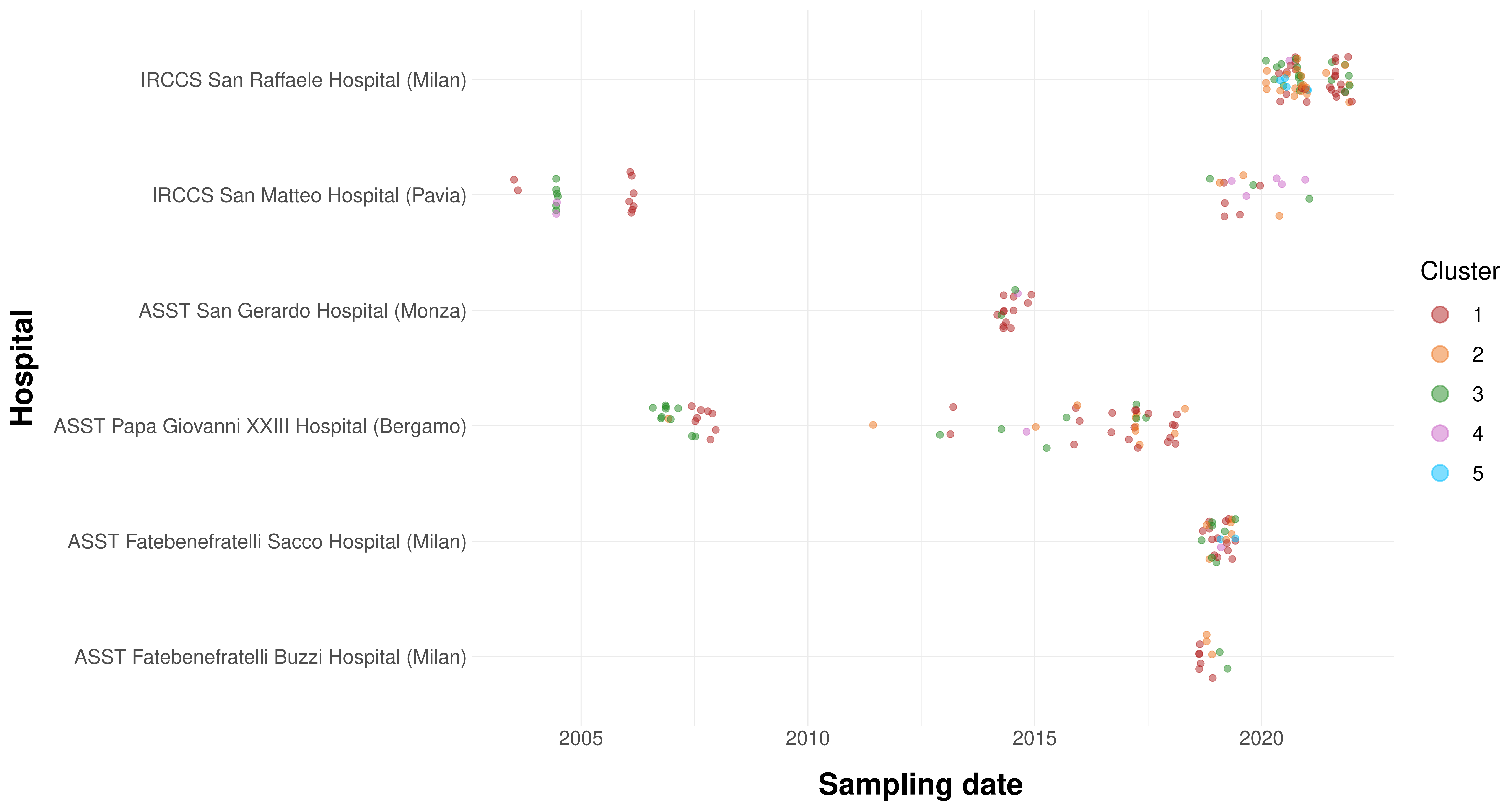

### Figure S6

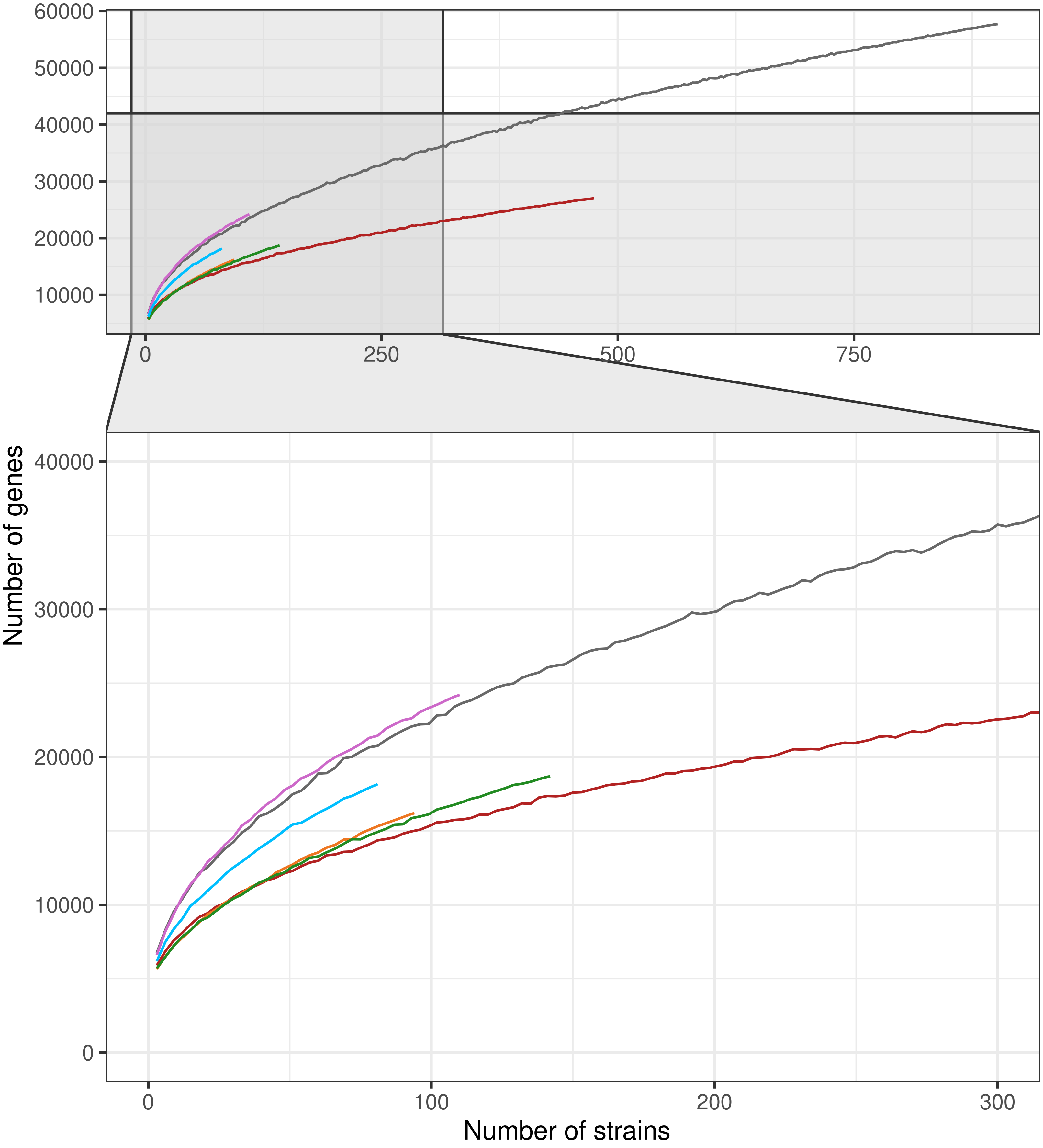

### Figure S7

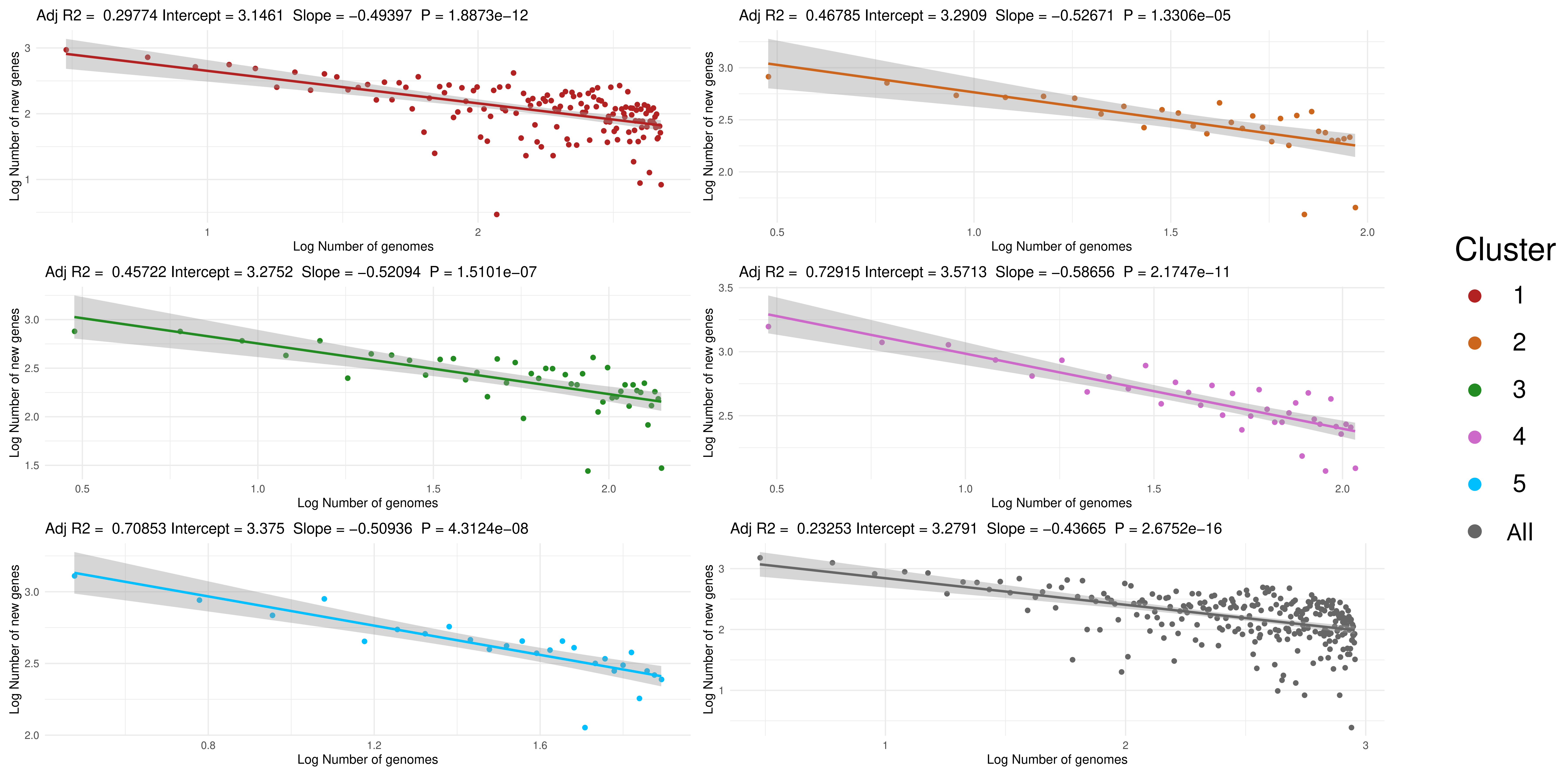

### Figure S8

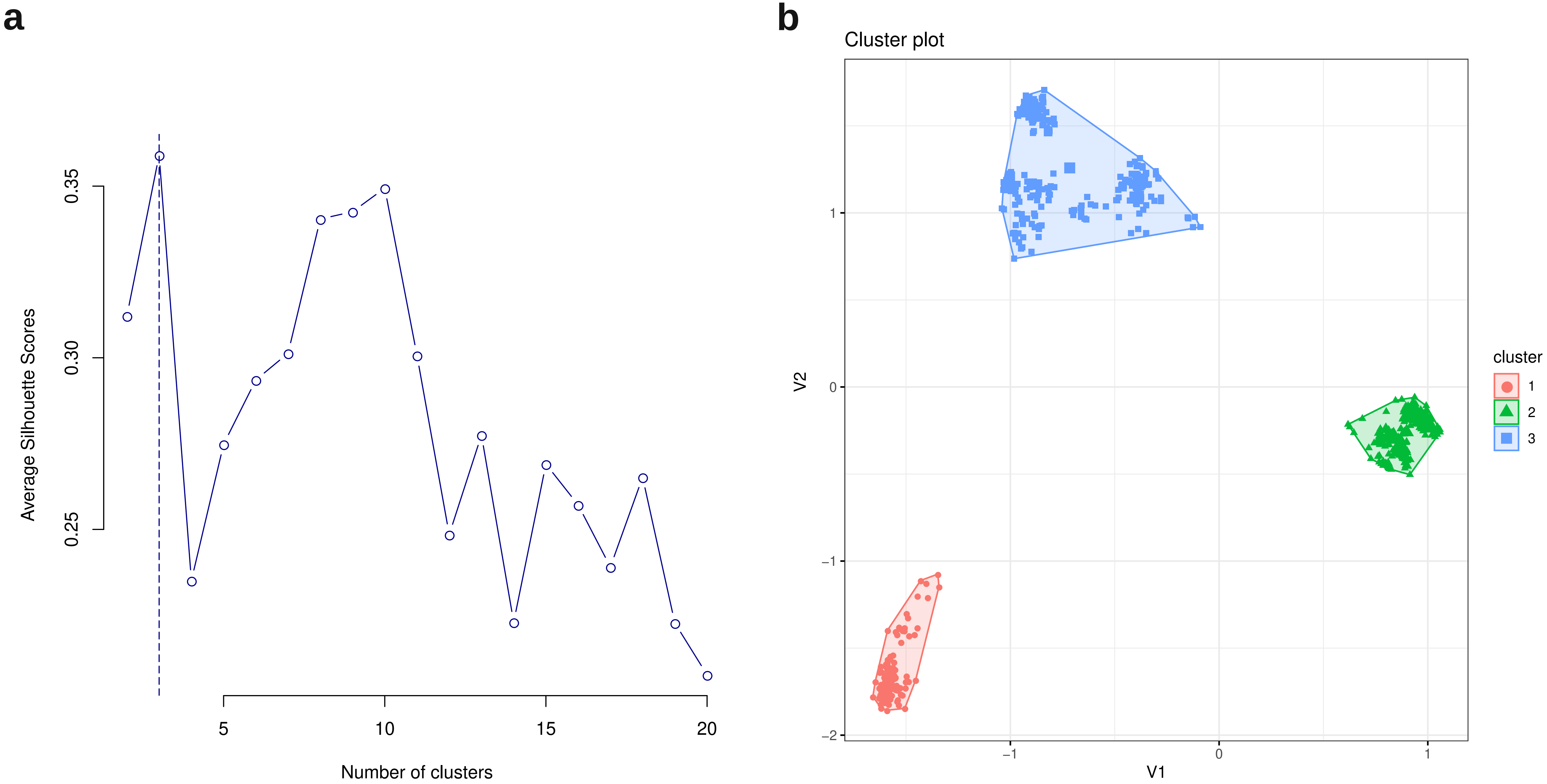

### Figure S9

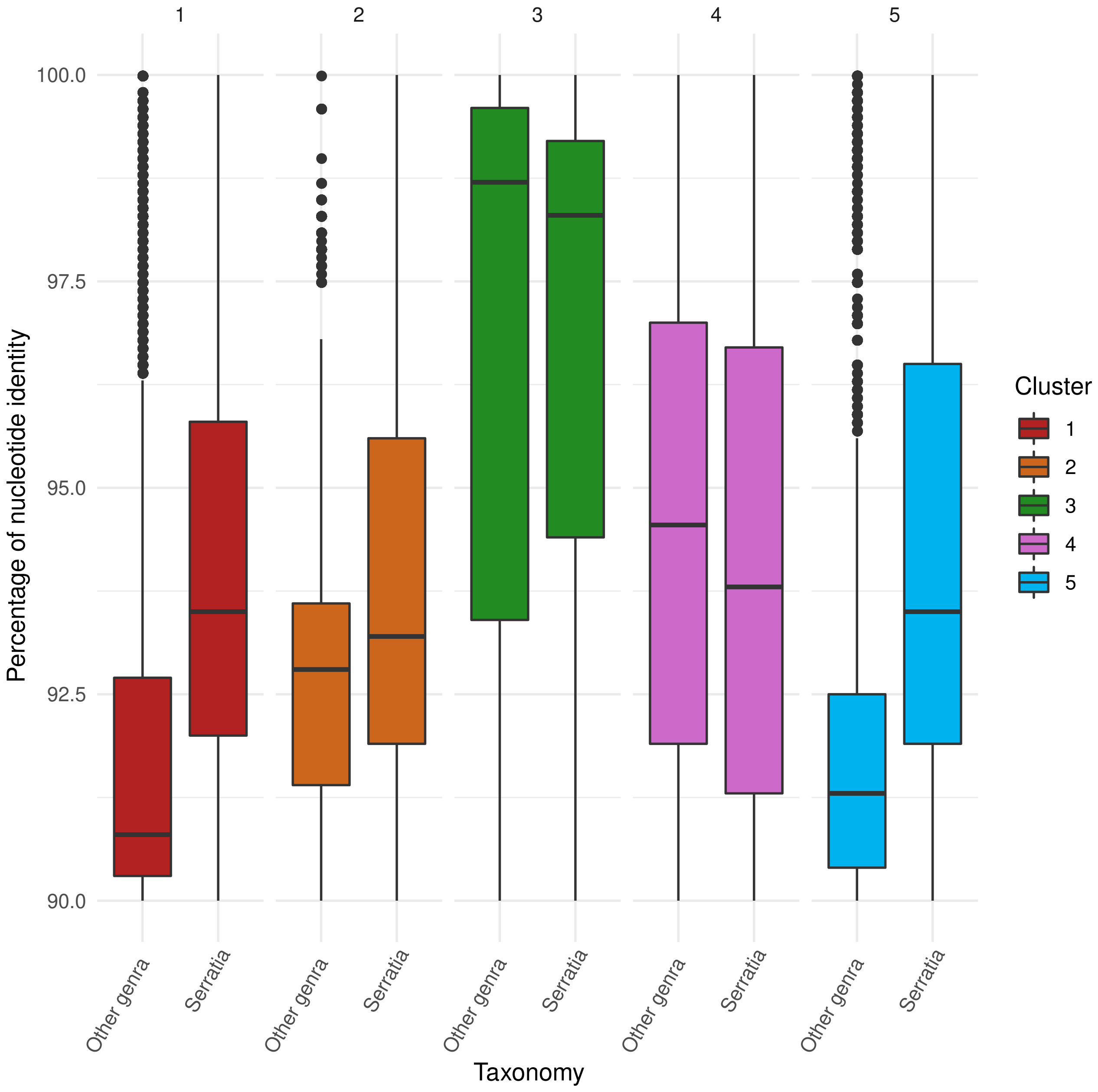

### Figure S10

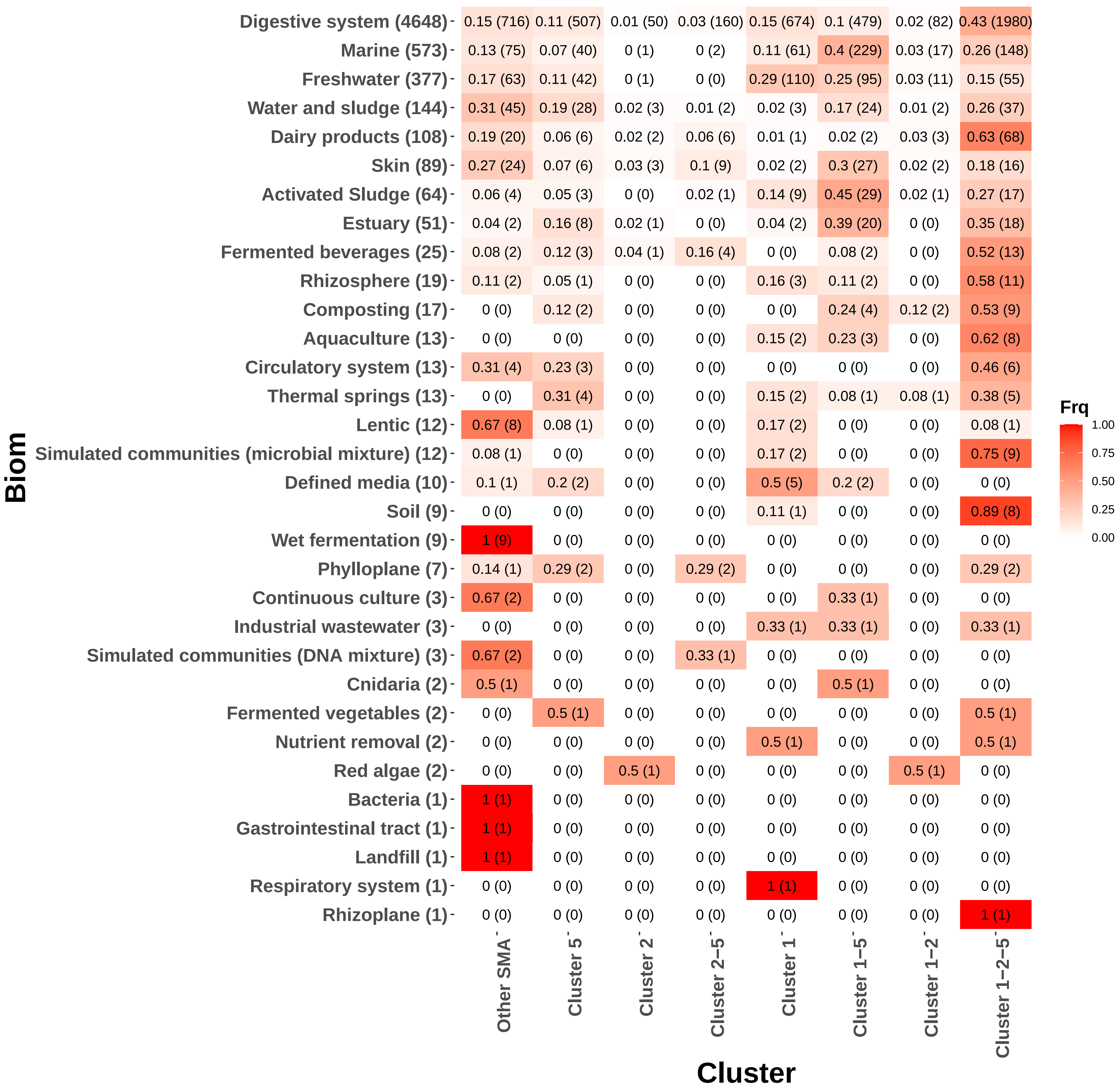

### Figure S11

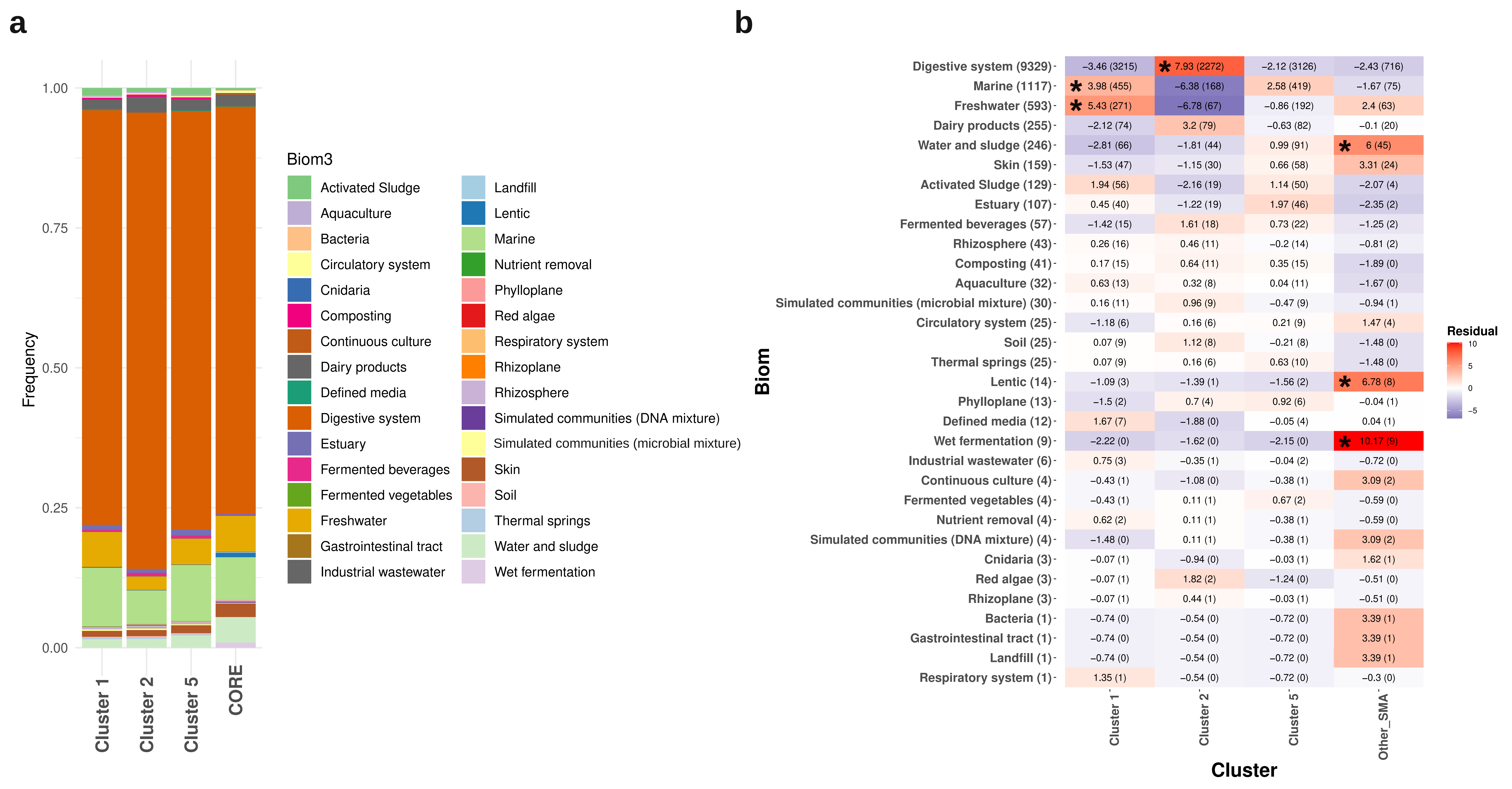

### Figure S12

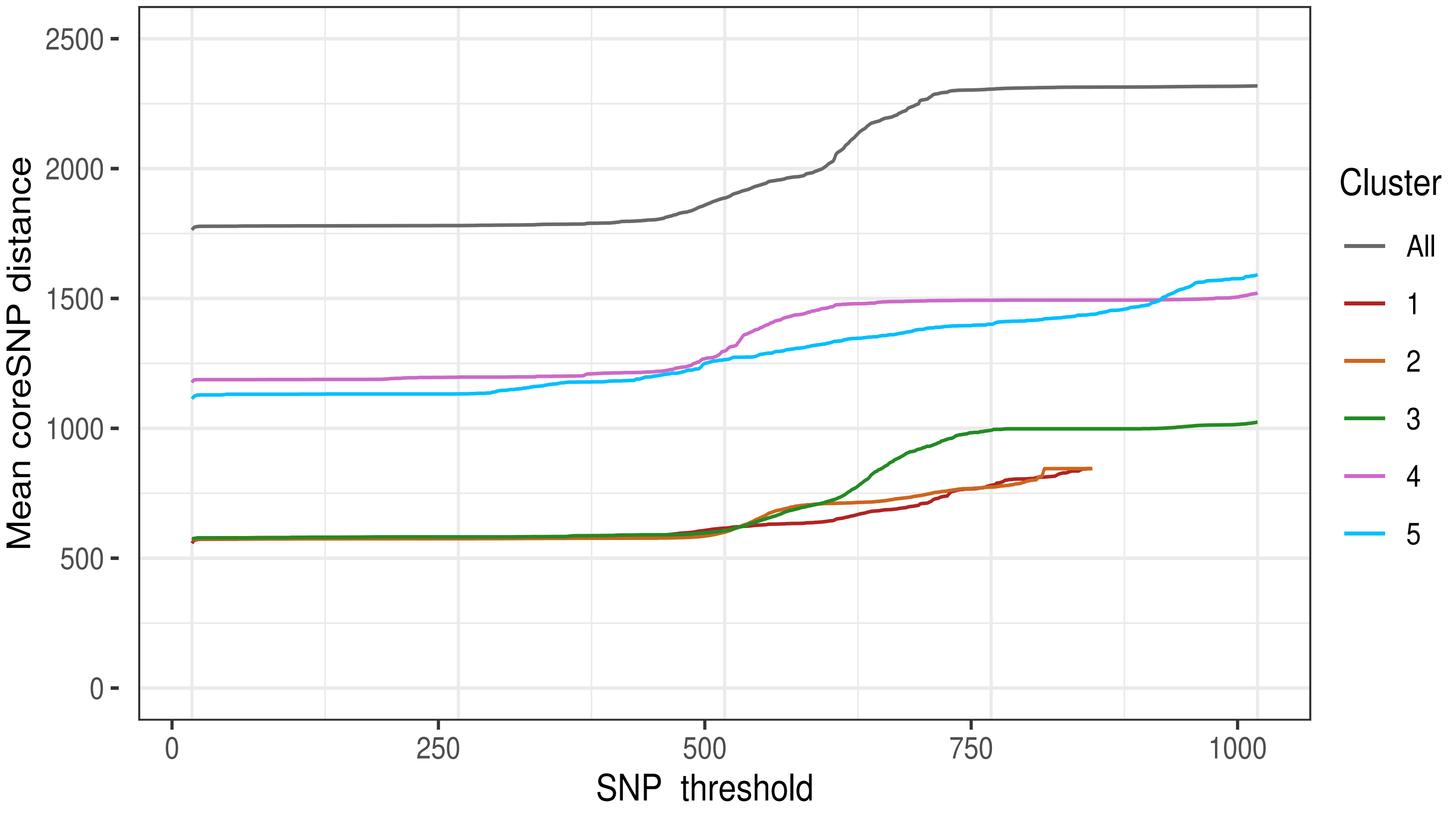

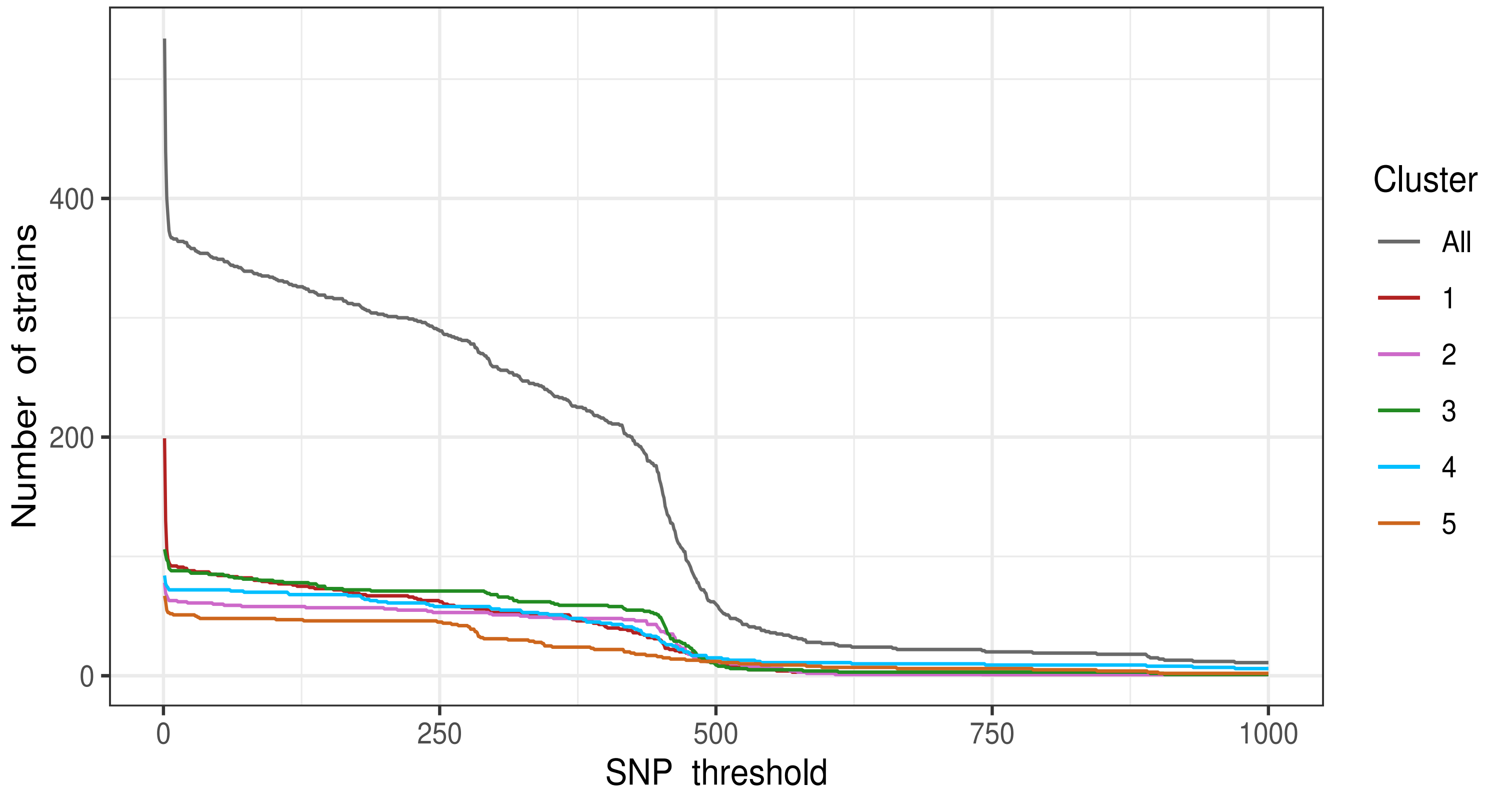

### Figure S13

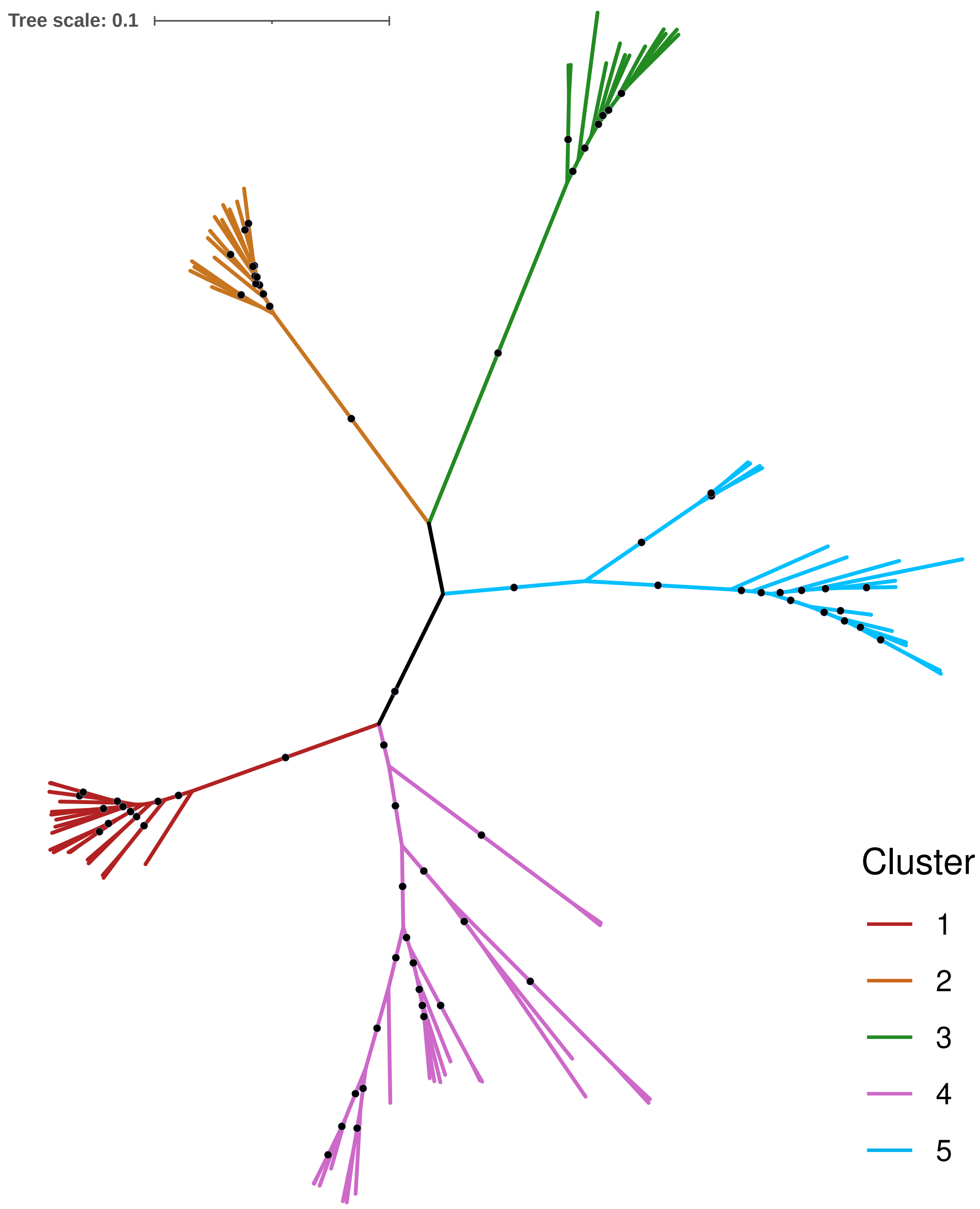

### Figure S14

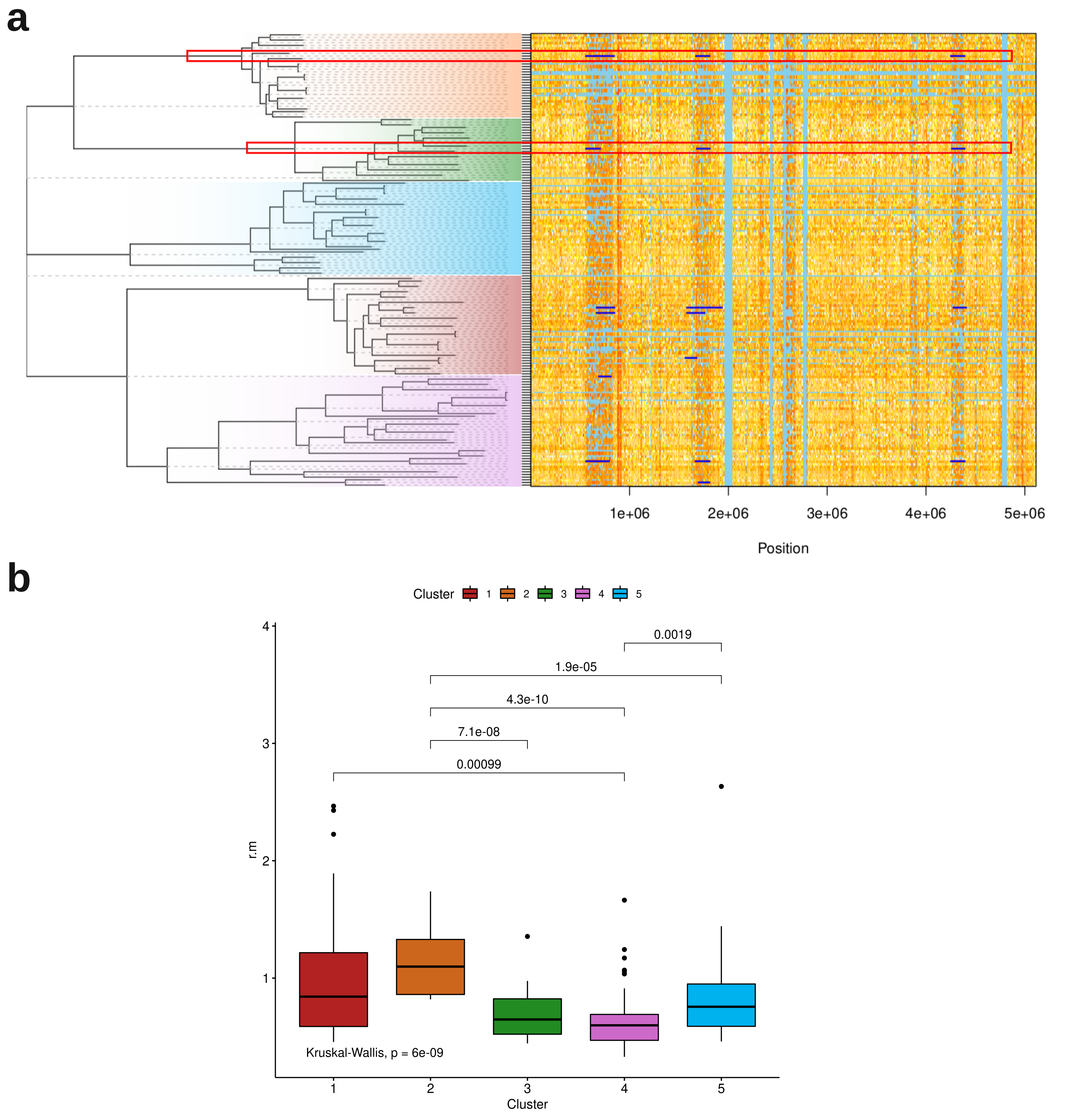

### Figure S15

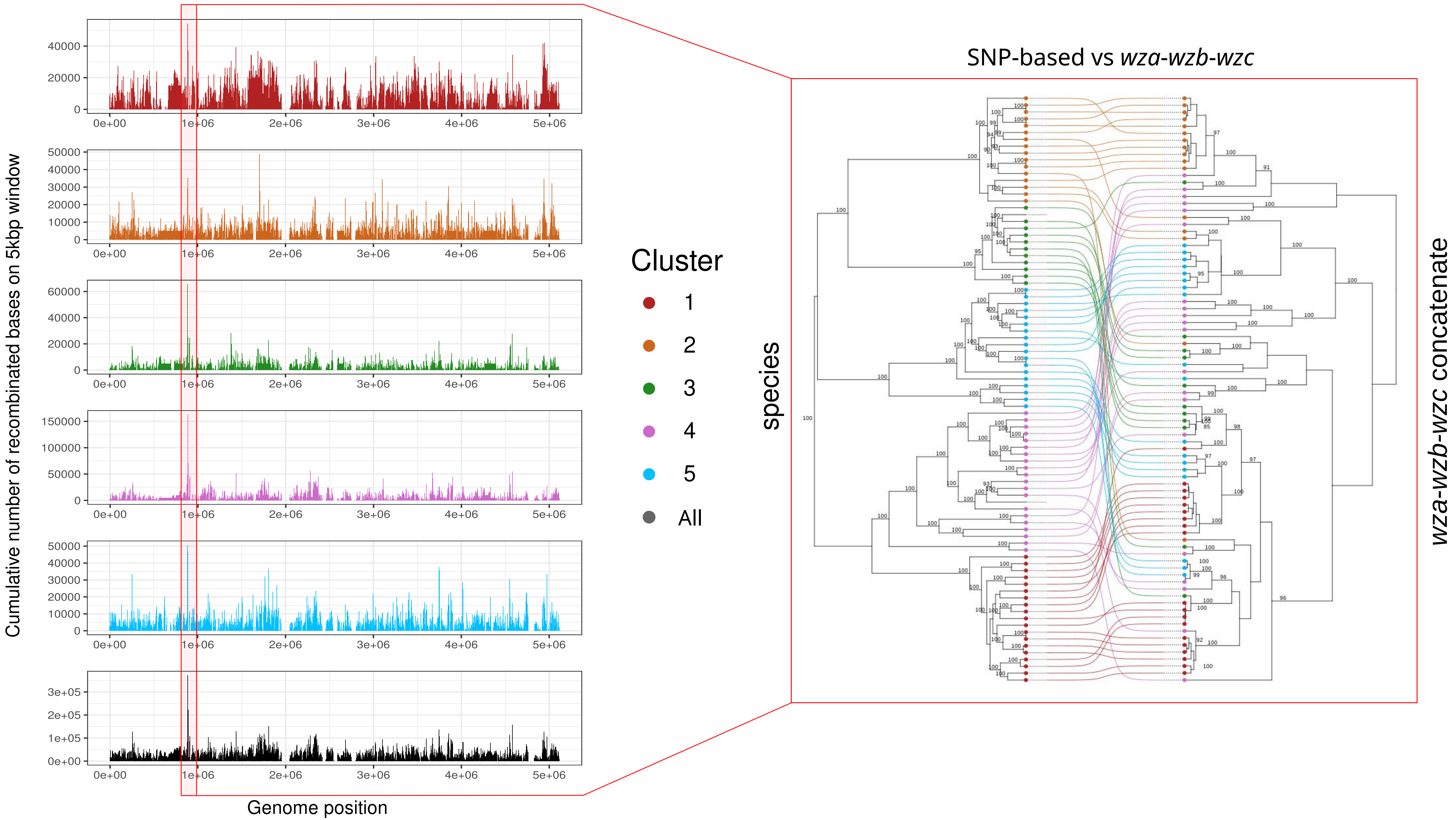
