## Supplemental material 1 for "Genetic barriers more than ecological adaptations shaped *Serratia marcescens* diversity"

**Supplementary material 1. Statistically significant P values in the comparison of genome size and GC content between *S. marcescens* clusters**

Genome size and GC content between strains of different clusters were compared by Mann-Whitney U-test with Holm post-hoc correction.

Genome size: Cluster 4 has a wider genome size in comparison to Cluster 1 (p = 1.8e-05, Mann-Whitney U test), Cluster 2 (p = 1.4e-07), Cluster 3 (p = 2.5e-10) and Cluster 5 (p = 2.2e-06). Cluster 1 genomes are also significantly larger than genomes in Cluster 3 (p = 0.00013, Mann-Whitney U test).

GC content: Cluster 1 has a markedly higher GC content than Cluster 2 (p < 2.22e-16, Mann-Whitney U test), Cluster 3 (p < 2.22e-16), Cluster 4 (p < 2.22e-16) and Cluster 5 (p = 3.8e-14). At the same time, Cluster 2 also has a lower GC content than Cluster 3 (p < 2.22e-16).
